# Development of ultra-low-input nanoRibo-seq enables quantification of translational control, revealing broad uORF translation by subtype-specific neurons

**DOI:** 10.1101/2022.04.05.487068

**Authors:** John E. Froberg, Omer Durak, Jeffrey D. Macklis

## Abstract

While increasingly powerful approaches enable investigation of transcription using small samples of RNA, approaches to investigate translational regulation in small populations of specific cell types, and/or (sub)-cellular contexts are lacking. Comprehensive investigation of mRNAs actively translated into proteins from ultra-low input material would provide important insight into molecular machinery and mechanisms underlying many cellular, developmental, and disease processes *in vivo*. Such investigations are limited by the large input required for current state-of-the-art Ribo-seq. Here, we present an optimized, ultra-low input “nanoRibo-seq” approach using 10^2^ – 10^3^-fold less input material than standard approaches, demonstrated here in subtype-specific neurons. nanoRibo-seq requires as few as 2.5K neurons, and exhibits rigorous quality control features: 1) strong enrichment for CDS versus UTRs and non-CDS; 2) narrow, distinct length distributions over CDS; 3) ribosome P-sites predominantly in-frame to annotated CDS; and 4) sufficient ribosome-protected fragment (RPF) coverage across thousands of mRNAs. As proof-of-concept, we calculate translation efficiencies from paired Ribo-seq and alkaline fragmented control libraries from “callosal projection neurons” (CPN), revealing divergence between mRNA abundance and RPF abundance for hundreds of genes. Intriguingly, we identify substantial translation of upstream ORFs in the 5’ UTRs of genes involved in axon guidance and synapse assembly. nanoRibo-seq enables previously inaccessible investigation of translational regulation by small, specific cell populations in normal or perturbed contexts.

## INTRODUCTION

Regulation of mRNA translation is crucial for establishing and maintaining appropriate protein abundances and sites of production, and thus for coordinating complex cellular functions. Dysregulated translation is an important mechanism underlying many human diseases, including ribosomopathies caused by mutations in the core ribosomal machinery[1], cancers driven by abnormalities in both signaling pathways regulating global levels of translation, as well as translation initiation and elongation factors themselves[2], and neurodevelopmental disorders such as Fragile X Syndrome[3] caused by mutations in RNA binding proteins that regulate protein synthesis. Despite the central importance of translational control in development and disease, it has been investigated far less extensively than transcriptional control in the same biological systems, due to both the relative simplicity of RNA-seq compared with more complicated biochemistry, and the dramatically larger input material requirements needed to determine translational output. New approaches are needed to enable analyses of translational output across distinct cell types, developmental stages, and genetic manipulations in specific cell populations isolated from appropriate *in vivo* contexts.

One such system with immense diversity of cell types, developmental dynamics of function, and critical relevance of genetic manipulations for cell type-specific disease is cerebral cortex neurogenesis and circuit formation-maintenance, in which translational regulation likely plays critical roles, but with immense gaps in knowledge due to inaccessibility of ultra-low input translational analyses. The mammalian cerebral cortex is organized into six layers, each layer with substantial neuronal diversity. Cortex differentiates from ventricular zone progenitors in an “inside-out” fashion, with deep-layer neurons born and differentiating earlier than superficial-layer neurons[4], thus with complex subtype-specific developmental dynamics. Cortical “projection” neurons are glutamatergic, excitatory neurons that extend exceptionally long axonal projections from their somata to distant targets 10^3^-10^5^ cell body diameters away, with diverse subtypes defined by soma location, and projection target(s)[4].

Callosal projection neurons (CPN) are an exemplar subtype. CPN project through the corpus callosum (largest axonal tract in the central nervous system) across the midline to homotypic targets in the contralateral hemisphere[5], and provide association and integration of functional areas in the two cortical hemispheres, underlying many cognitive, behavioral, and associative sensorimotor functions [6, 7]. Approximately 80% of CPN are “late-born” superficial layer (II/III) neurons, with distinct populations of CPN in deeper layers. Beyond their subtype-specificity of axonal targeting and function, and their highly orchestrated developmental dynamics, numerous human genetic variants and mutations disrupt CPN circuitry in humans and mice, causing distinct, subtype-specific behavioral dysfunction. Both transcriptional regulation and translational regulation likely play combinatorial and distinct roles in these circuits, functions, and dysfunctions.

Post-transcriptional control has the potential to restrict protein expression to proper developmental times, cell/neuron subtypes, and subcellular locations, highlighting the likely complex interplay between transcriptional and translational regulation. For example, cortical progenitors and newborn neurons during early differentiation often transiently co-express mRNAs coding for subtype identity-specifying transcription factors (TFs) of multiple subtypes[8–13], but do not produce these TF proteins[8–11]. In addition, shifts in expression of RNA binding proteins including Pum1/2[9, 11], 4E-T[9, 10], HuR[14], HuD[15], Ire1α[16], Elav4, and Celf1[17], and of ribosomal proteins Rpl7 and Rpl10[18], control translation of select mRNAs in progenitors while neurogenesis shifts from production of deep-layer neurons to superficial-layer neurons such as CPN[9-11, 14-18]. The reported global decrease in ribosome biogenesis and abundance from early to late cortical neurogenesis might also favor translation of select sets of genes at progressive developmental stages[11, 18, 19]. These studies, while mostly from bulk neocortex without isolation of particular progenitor classes or distinct neuron subtypes, also suggest an important role for translational regulation in fine-tuning output of developmental programs that define subtype identity. Importantly, much less is known about translational regulation by distinct cortical subtypes, especially at later developmental stages as their long-range projections and precise circuitry develop. Investigating translational output by specific subtypes requires development of new approaches to analyze translation from the extremely limited material that can be obtained from purified subtype-specific neurons.

We adapted Ribo-seq (also termed ribosome profiling) to the ultra-low quantities of material practicably obtainable from distinct cortical projection neuron subtypes, enabling investigation of subtype- and stage-specific translational regulation. Ribo-seq provides global measurement of translational output[20, 21], but standardly requires input material from ∼10^6-7^ cells. For Ribo-seq analysis, a cell lysate is digested with an RNase under controlled conditions to enrich for “ribosome protected footprints” (RPFs), short fragments of mRNAs protected from digestion by the bulk of the ribosome complex. Fragments in the expected RPF size range are size-selected and sequenced. Ribo-seq offers the advantageous capability that one read corresponds to one translating ribosomal complex, and RPF abundance can be compared with global mRNA abundance to infer the “translational efficiency” (T.E.)[21] for each mRNA. Two features of Ribo-seq provide robust “quality controls” to assess whether the bulk of reads correspond to bona fide translating ribosome footprints, rather than other processes. First, RPFs have characteristic length distributions (typically ∼30-nt in mammalian systems)[22–24], and the putative position of a translating ribosome P-site can be inferred from the RPF size distribution with single nucleotide precision[20, 21]. Second, P-sites predominantly match the positions of in-frame codons, leading to a pronounced 3-nt periodic signal[20, 21].

Nucleotide-level resolution of Ribo-seq data also enables robust analysis of complex phenomena such as translation of upstream ORFs (uORFs)[21, 25, 26], elongation[27] and termination[28] dynamics, even co-translational translocation of secreted proteins into the endoplasmic reticulum[29]. Ribo-seq is also uniquely powerful for detecting novel translated ORFs, often encoding small peptides[30–32], which can have substantial clinical implications, since some of these are neoantigens in malignant tumors[33], possibly enabling targeted cancer vaccines[33].

The main limitation of Ribo-seq is that standard protocols generally require very large amounts of material (10^6-7^ mammalian cells; 10-100 ug RNA)[21], far in excess of what can be obtained from relatively rare or otherwise relatively inaccessible cell types of interest. Lower-input Ribo-seq methods have been reported, e.g. examining translation in hippocampal slices[34], and using *in vitro* axon outgrowth systems[35], but these approaches still require microgram-scale quantities of RNA. A recent study reported Ribo-seq from single cells[36], but this approach measured only RPF abundance without comparator mRNA abundance for T.E. measurements, had only modest enrichment for in-frame P-sites, and requires highly specialized robotics and microfluidics. A generalizable ultra-low-input Ribo-seq approach would enable powerful new investigations of type-specific, developmentally stage-specific, and context-specific cell populations.

To enable cell type-specific and potentially subcellular analyses of translational regulation across a wide variety of biological systems, we present here an optimized ultra-low-input Ribo-seq approach appropriately scaled to facilitate Ribo-seq with nanogram-scale RNA inputs (nanoRibo-seq). We present results from cortical projection neurons as an exemplar proof of concept (Figure1). We first present a general framework for optimizing Ribo-seq using just 30 ng of RNA as starting material (Figure 1A), leveraging the fact that optimal Ribo-seq libraries should exhibit the following features (Figure 1-figure supplement 1): 1) enrichment of RPFs over coding sequence (CDS) compared to untranslated regions; 2) distinct fragment length distributions over CDS versus non-CDSs; 3) 3-nt, in-frame periodicity of inferred ribosome P-sites over CDS.

**Figure 1:**
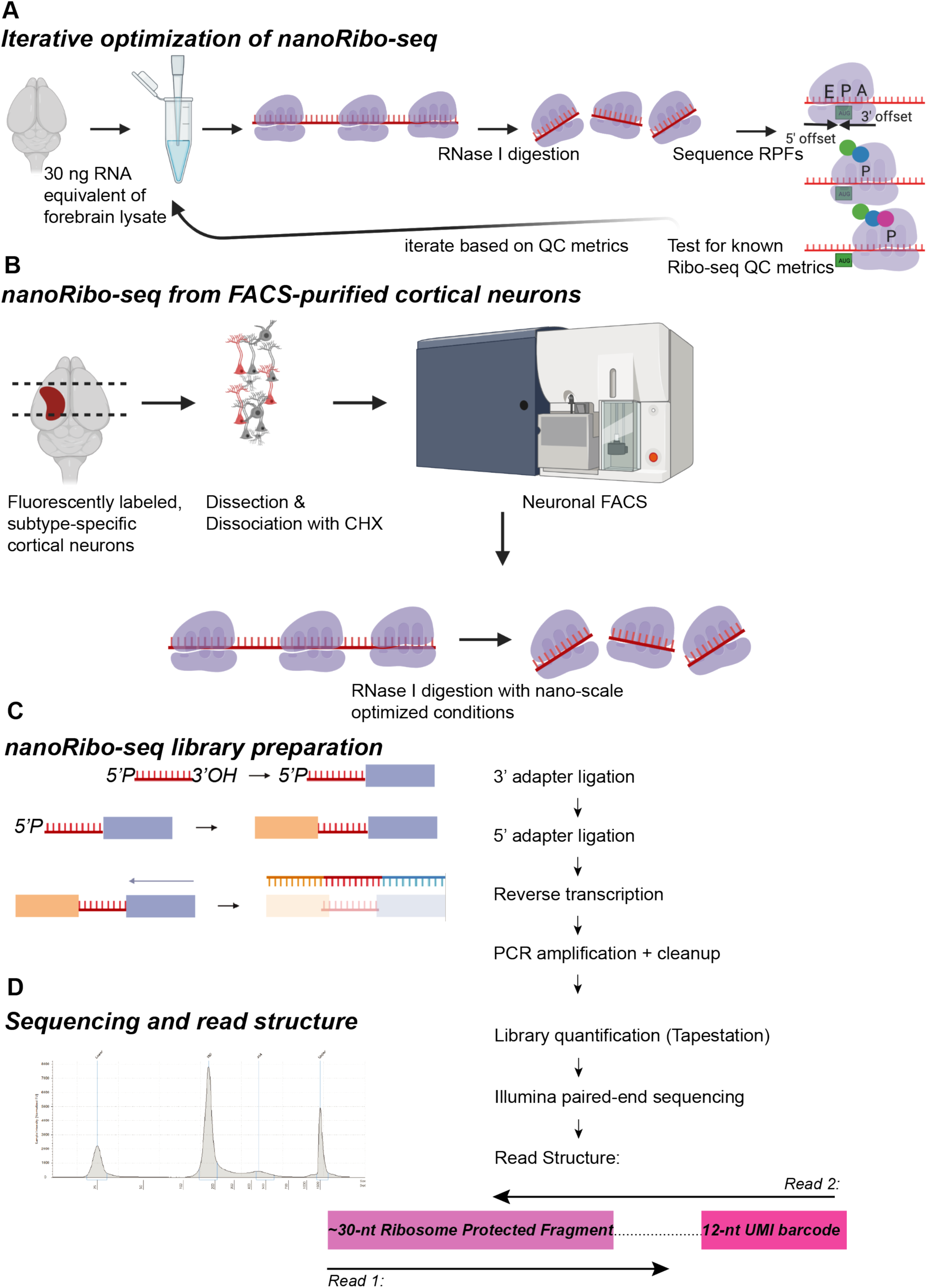
Schematic of nanoRibo-seq approach to investigate translational regulation by defined neuronal subtypes or subpopulations. A) Iterative optimization of nanoRibo-seq: We tested RNase I digestion conditions using 30 ng of forebrain lysate, isolated and sequenced RPFs, and tested for the presence of well-known ribosome profiling QC metrics 1) enrichment over CDS, 2) distinct length distributions over CDS, and 3) enrichment in-frame P-sites). We iteratively repeated these nano-scale RNase I digestions until the QC metrics were optimized. B) Generalized workflow for nanoRibo-seq from labeled, FACS-purified cortical neurons. C) Workflow for conversion of nano-scale quantities of small RNA into RNA-seq libraries using the QiA-seq miRNA library kit, an RNA ligation-based method. D) Library preparation results and read structure: Resulting libraries have a sharp size distribution centered at ∼180-bp, consistent with a roughly 30 bp insert between the two sequencing adapters. This results in a read structure in which Read 1 sequences the ∼30 nt RPF, and Read 2 sequences the 12-nt UMI barcode, enabling identification and removal of PCR duplicates before quality control and/or quantitative analyses.

We report successful, high-depth coverage nanoRibo-seq in CPN, and in an even rarer population of cortical projection neurons that project to the brainstem and spinal cord (subcerebral projection neurons; SCPN), using as few as 2500 FACS-purified neurons. We also optimized preparation of alkaline fragmented control libraries for measuring mRNA abundance from this same ultra-low input. Combining the two, we generated paired nanoRibo-seq (RP), and alkaline fragmented (AF) samples using FACS-purified CPN. These methods enable detection of thousands of genes in each nanoRibo-seq experiment, along with calculation of translation efficiency (T.E.)-– the log ratio of RPF abundance from RP experiments, and mRNA abundance from AF experiments (log_2_(RP/AF))-– for thousands of detected transcripts. The results reveal that, in the exemplar CPN, approximately 10% of genes display a significant difference between RPF and mRNA abundance, indicating potential translational regulation of hundreds of genes. Strikingly, these unique data identify that a select set of short upstream open reading frames (uORFs) are actively translated by CPN. uORFs are initiated within mRNA 5’ UTRs, and are often strongly inhibitory for translation of the main CDS, though are also typically poorly translated under constitutive conditions[37]. Our data reveal that genes with translated uORFs have lower T.E. than genes lacking translated uORFs. Intriguingly, transcripts coding for genes involved in synapse formation, synapse maintenance, and neuronal cell adhesion are enriched among the mRNAs containing translated uORFs. Thus, nanoRibo-seq enables investigation of translational regulation using ultra-low input quantities from subtype-specific and otherwise rare cell types, with results indicating substantial translational regulation in CPN. nanoRibo-seq can be applied using only a few thousand cells, uniquely enabling study of translational regulation by a wide range of cellular, tissue, and developmentally dynamic systems.

## RESULTS

### nanoRibo-seq yields rigorous results with as little as 30 ng RNA input

To investigate the feasibility of nanoRibo-seq in a relatively simple system with ultra-low input, we harvested neonatal mouse brains, lysed them in 1X polysome buffer (with cycloheximide to inhibit translational elongation), extracted RNA, measured its concentration, and calculated the approximate volume of brain lysate equivalent to 30 ng extracted RNA. We diluted this volume in polysome buffer (to approximate volumes obtained after FACS, with additional dilution to minimize effects of sheath fluid and neuronal disassociation buffer), and digested with RNase I. We selected the concentration of RNase I “[RNase I]” that produced an apparent RPF of approximately 30-nt (Figure 2A). In parallel, we performed “bulk” digestion of undiluted brain lysate using RNase I digestion conditions similar to the original mammalian Ribo-seq method[21]. We also digested purified brain lysate total RNA with RNase I for comparison to Ribo-seq libraries (Figure 2-figure supplement 1).

**Figure 2:**
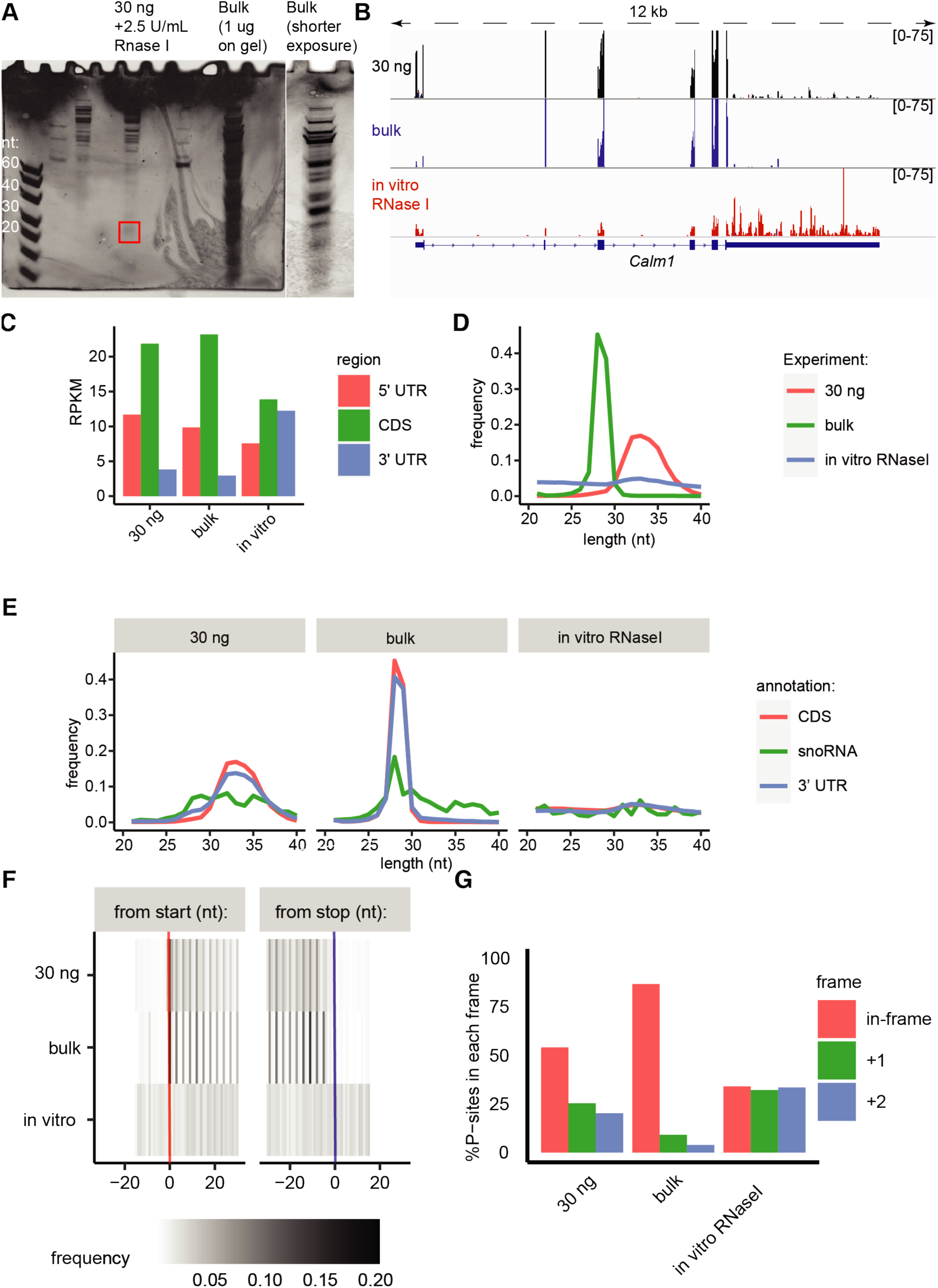
Ribo-seq from neonatal brain lysates with either ultra-low (30 ng) or bulk quantities of material. A) RNase I digestion patterns with either brain lysate volume equivalent to 30 ng RNA + 2.5 U/mL RNase I or “bulk” conditions: 120 ug RNA+1500 U RNase I. The red boxed region indicates putative ribosome-protected fragments. B) Read coverage over the *Calm1* gene in 30 ng (top, black) or bulk (middle, blue) Ribo-seq libraries, or in the *in vitro* RNase I-digested total RNA library (bottom, red). C) RPKM coverage (reads/kilobase)/(million uniquely mapped) over CDS (5’ UTR=red, CDS=green, 3’ UTR=blue). D) Fragment length distributions over CDS (red=30 ng, green=bulk, blue=in vitro digested). E) Fragment length distributions over CDS (red), snoRNAs (green), 3’ UTRs (blue), and for 30 ng library (left), bulk library (center), or in vitro digested (right). F) Heatmap of P-site relative frequency as a function of position from the start (red) or stop codon (blue). G) % of P-sites mapping to each reading frame (in-frame=red, +1=green, +2=blue).

To convert these short RNA fragments into sequencing libraries, we employed the QiA-Seq miRNA library kit, designed for generation of small RNA libraries with as little as 1 ng RNA, and we used paired-end sequencing. The QiA-Seq miRNA library kit introduces a 12-nt unique molecular identifier (UMI) sequenced at the start of Read 2 in Illumina paired-end sequencing (Figure 1D), thus enabling PCR duplicates to be collapsed and removed before all analysis. Importantly, PCR duplicates frequently confound quantification in low-input genomics approaches that do not allow for identification and removal of PCR duplicates using UMIs. Following duplicate removal, we analyzed the libraries for characteristics that indicate capture of elongating ribosome protected footprints: 1) higher coverage in CDS (CDS) than UTRs, especially 3’ UTRs; 2) distinct read length distributions in CDS versus ncRNA and, 3) a 3-nt, periodic enrichment of inferred P-sites in-frame with the annotated CDS (Figure 1-figure supplement 1).

Both the bulk and 30 ng Ribo-seq libraries, but not the RNase I digested total RNA control library exhibit higher coverage over CDS than both 5’ and 3’ UTRs (Figure 2B & C). The Ribo-seq libraries exhibit tight, unimodal fragment length distributions (consistent with ribosome protection), but RNase I digested total RNA does not show any specific preferred length (consistent with random digestion) (Figure 2D). The Ribo-seq libraries also show narrower length distributions over CDS compared to 3’ UTRs and snoRNAs (Figure 2E). This is important since structured RNAs or tightly bound RNA binding proteins might also protect RNA from digestion, but are unlikely to protect RNA in exactly the same way as the ribosome, leading to distinct length distributions between CDS and non-coding RNA[11].

Another hallmark of successful Ribo-seq is the distinct enrichment of P-sites in-frame with the dominant CDS every 3-nt. We used the RiboWaltz R package[38] to infer P-sites. There was a clear 3-nt, in-frame periodicity to inferred ribosomal P-sites in both the bulk and 30 ng libraries around both the start and stop codons (Figure 2F), and enrichment of in-frame P-sites versus out-of-frame P-sites (Figure 2G). This enrichment of in-frame P-sites strongly indicates that the ultra-low input nanoRibo-seq approach captures translating ribosomes, since ribosomes move codon-to-codon in 3-nt steps, and few other biological processes can produce such a distinct 3-nt periodicity, in-frame, specifically within CDS. Together, these initial experiments indicate that Ribo-seq is feasible with very small quantities of brain lysate material, since both the 30 ng and bulk libraries exhibit the hallmarks of translating ribosomes.

### RNase I titration reveals optimal digestion conditions for nanoRibo-seq

Of note, the 30 ng library displayed a longer, broader length distribution than the bulk library (Figure 2D), slightly longer than the ∼29-31 nt reported in most mammalian Ribo-seq studies[21, 22, 34]. It also displayed a weaker 3-nt periodicity, and weaker enrichment for in-frame P-sites compared to the bulk sample (Figure 2F & G), which might arise due to decreased accuracy in inferred P-site positions from a longer, broader length distribution.

We hypothesized that more limited RNA digestion underlies the longer and broader length distribution in the 30 ng library. Thus, we titrated [RNase I], to investigate whether there exist low-input conditions producing tighter size distributions. We tested the same [RNase I] as the original experiment (2.5 U/mL), and a range of increasing [RNase I] (12.5 U/mL, 62.5 U/mL, 312.5 U/mL) (Figure 3A). All four enzyme concentrations yield enrichment of CDS over 5’ and 3’ UTR (Figure 3B & C), with the strongest enrichment at 12.5 U/mL. We found that increasing [RNase I] shortens and tightens the fragment length distribution over CDS, and the higher concentrations yield footprint lengths in line with previous observations (∼29-31 nt) (Figure 3D). All digestion conditions yield distinct length distributions over CDS compared to 3’ UTRs (Figure 3E). All libraries also displayed an in-frame, 3-nt P-site periodicity (Figure 3F), and enrichment of in-frame P-sites (Figure 3G). The 12.5 U/mL and 62.5 U/mL conditions displayed the strongest periodicity, and the highest percentage of in-frame P-sites. Together, these results demonstrate that a wide range of ultra-low input digestion conditions generate libraries meeting the three rigorous and stringent quality control metrics for Ribo-seq, matching those previously reported using inputs 2-3 orders of magnitude higher than those used here[34].

**Figure 3:**
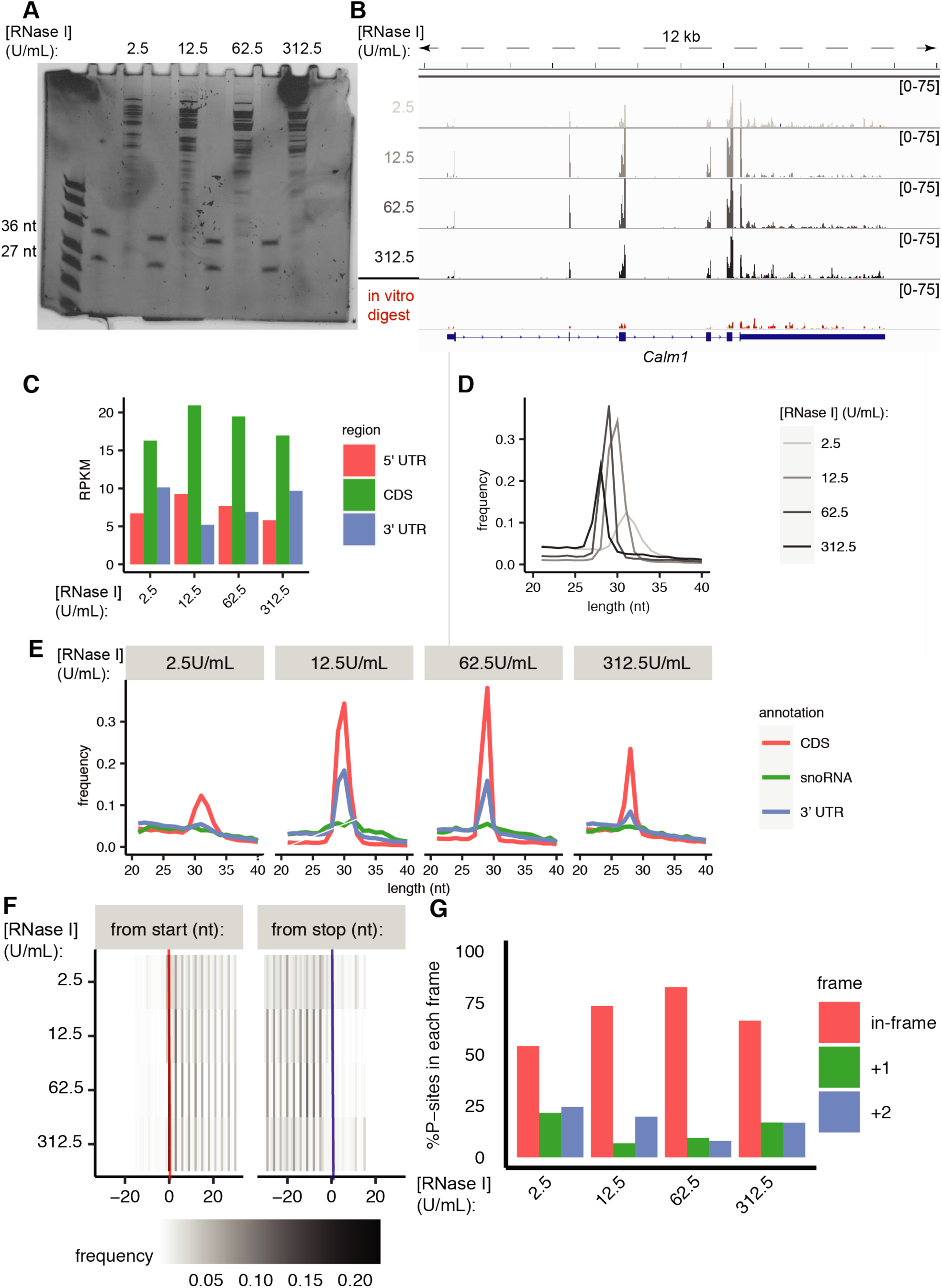
RNase I titration against 30 ng brain lysate. A) RNase I digestion patterns at RNase I concentration=2.5, 12.5, 62.5, 312.5 U/mL. B) Read coverage over the *Calm1* gene in Ribo-seq libraries (light to dark grey with increasing [RNase I]) vs. *in vitro* RNase I digested total RNA (red). C) RPKM coverage (reads/kilobase)/(million uniquely mapped) over CDS (5’ UTR=red, CDS=green, 3’ UTR=blue). D) Fragment length distributions over CDS (light to dark grey with increasing [RNase I]). E) Fragment length distributions over CDS (red), snoRNAs (green), 3’ UTR (blue) at four distinct RNase I concentrations. F) Heatmap of P-site signal counts as a function of position from the start (red) or stop codon (blue). G) % of P-sites mapping to each reading frame (in-frame=red, +1=green, +2=blue).

### Alkaline fragmentation produces superior low input total RNA control libraries compared with RNase I digestion

An important comparator for Ribo-seq experiments is a fragmented total RNA library for simultaneous measurement of RPF abundance and mRNA abundance, ideally generated with library methods similar to those employed for the Ribo-seq library. We tested three strategies for generating fragmented total RNA control libraries with small quantities of RNA isolated from early postnatal forebrain lysate (Figure 2-figure supplement 1). We first tested RNase I digestion of purified RNA (Figure 2-figure supplement 1A & B). Since rRNA comprises >90% of total RNA samples, we also tested two rRNA removal strategies to enrich fragmented control libraries for mRNA. We performed rRNA removal with the NEB human/mouse rRNA Removal Kit after gel purification (“G->R method”), but found that a large amount of short hybridization probe sequences contaminates the libraries (Figure 2-figure supplement 1C). Thus, we also tested rRNA removal before gel purification to remove the short probes (“R->G method”), since most run below the expected RPF size. Although RNase I digestion would enable complete matching of library preparation protocols between the fragmented control and Ribo-seq libraries, we found that RNase I produced broad fragment size distributions, and had a very narrow concentration window between producing desired ∼20 to 60-nt fragment sizes and complete digestion (Figure 2-figure supplement 1B).

Some Ribo-seq approaches instead utilize alkaline fragmentation for the control libraries[20, 21]. We produced an alkaline fragmented library from 7.5 ng purified RNA–without rRNA removal, both for simplicity, and to optimally match the Ribo-seq procedure itself, which also lacks rRNA removal (Figure 2-figure supplement 1D). We found that alkaline fragmentation produces tighter size distributions, especially those within the desired 20 to 60-nt window (Figure 2-figure supplement 1E). We also comparatively examined the mapping characteristics of the alkaline fragmented libraries and those generated using the two RNase I methods. Over half of reads in the “RNase I G->R method” were too short for alignment, because they are short probe sequences (Figure 2-figure supplement 1F). The “RNase I R->G method” had fewer reads too short for alignment, but actually fewer mRNA-aligned reads (Figure 2-figure supplement 1F & G). Though the alkaline fragmentation method generated a high proportion of rRNA-aligning reads (Figure 2-figure supplement 1F), approximately 5% of reads aligned to mRNA, roughly comparable to the fraction of reads aligning to mRNA in the Ribo-seq libraries themselves (Figure 2-figure supplement 1G). Importantly, we are able to obtain deep coverage over mRNA despite such high rRNA mapping rates because of deep sequencing (30-50M reads per library). Given the simplicity of alkaline fragmentation, the resultant tight size distributions, and the fact that the Ribo-seq libraries also omit an rRNA depletion step, alkaline fragmentation appears to be superior for producing low-input fragmented total RNA control libraries.

### High quality nanoRibo-seq from FACS-purified subtype-specific neurons

We next investigated whether nanoRibo-seq is amenable to sorted, subtype-specific CPN (and a rarer cortical neuron subtype, SCPN) under a variety of digestion and sorting conditions. We generated test libraries using a range of cell numbers, [RNase I], and both distinct subtypes (Figure 4A). We first dissected and dissociated P3 cortex labeled via E14.5 *in utero* electroporation (which introduces fluorophore-encoding plasmids into cortical progenitors), to label superficial layer CPN (with cycloheximide in all buffers from brain harvest onwards), and sorted 100K CPN (Figure 4-figure supplement 1A). To test [RNase I] concentrations, we split these 100K CPN into two sets of 50K CPN. One set was digested with 12.5 U/mL RNase I, found to be well within the optimal range in the titration experiment above using 30 ng brain lysate, and the other set with 2.5 U/mL to maintain a constant digestion condition across all experiments (Figure 4-figure supplement 1A). We also performed a second FACS to investigate with even lower levels of input material, collecting 17K P3 CPN, and digested with 12.5 U/mL (Figure 4-figure supplement 1A). In addition to these experiments using CPN, using conditions previously yielding successful Ribo-seq libraries, we tested the approach using a second subtype, the rarer subcerebral projection neurons (SCPN) without cycloheximide. Though many ribosome footprints in this initial SCPN sample likely arise from *ex vivo* protein synthesis, this enabled comparison of global features of interest to evaluate nanoRibo-seq in distinct subtypes. Thus, we sorted 50K SCPN into polysome buffer containing cycloheximide, and digested with 12.5 U/mL RNase I (Figure 4-figure supplement 1B).

**Figure 4:**
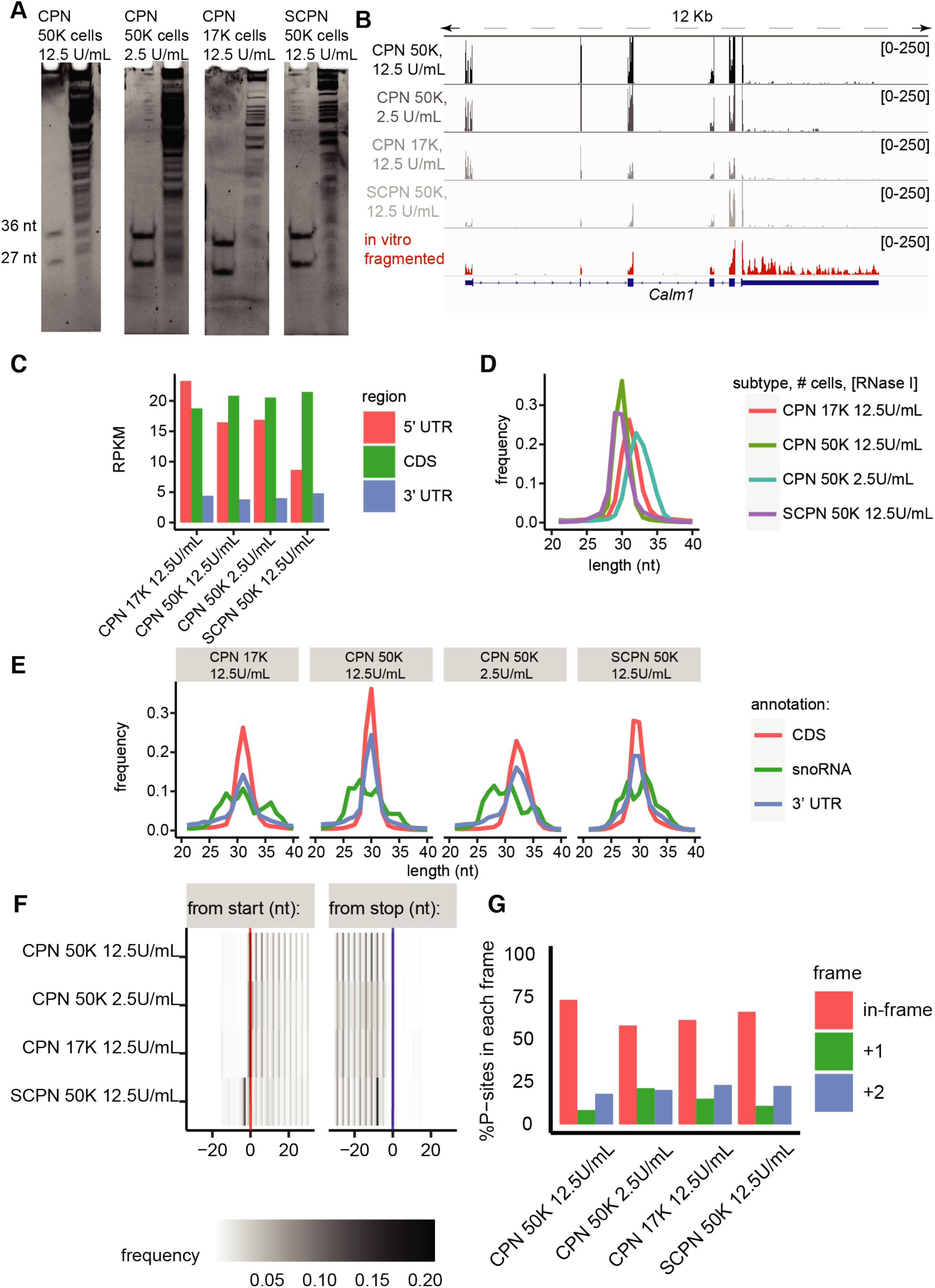
nanoRibo-seq quality control metrics from several thousand FACS-sorted CPN and SCPN. A) RNase I digestion patterns from sorted CPN and SCPN; number of cells and RNase I concentration indicated above lane. B) Read coverage over the *Calm1* gene from CPN and SCPN Ribo-seq libraries (shades of grey) versus *in vitro*, alkaline fragmented RNA. C) RPKM coverage (reads/kilobase)/(million uniquely mapped) over CDS (5’ UTR=red, CDS=green, 3’ UTR=blue). D) Fragment length distributions over CDS. E) Fragment length distributions over CDS (red), snoRNAs (green), 3’ UTR (blue). F) Heatmap of P-site signal counts as a function of position from the start (red) or stop codon (blue). G) % of P-sites mapping to each reading frame (in-frame=red, +1=green, +2=blue).

All four CPN and SCPN libraries exhibited quality control indicators of successful and high quality ribosome profiling. Each of the libraries displayed several fold higher enrichment of CDS over 3’ UTRs (Figure 4B & C). The footprint length distributions over CDS were observed to depend on both enzyme concentration and cell number (Figure 4D); the 12.5 U/mL CPN sample displayed a shorter and narrower length distribution than the 2.5 U/mL CPN sample. Interestingly, the 12.5 U/mL CPN and SCPN samples displayed the most similar length distributions, despite the fact that they are isolated from two distinct cortical projection neuron subtypes, labeled using different strategies, sorted on different days, and subjected to different translation inhibition regimes. The 17K neurons, 12.5 U/mL CPN library displayed a slightly wider and broader length distribution than the 50K neurons, 12.5 U/mL library. This suggests that cell concentration affects consistent digestion, since both digestions used the same enzyme concentration in the same volume, thus the concentration of neurons in the 50K CPN sample was ∼3X higher than the 17K sample. Together, these results indicate that digestion depends most critically on [RNase I] and cell concentration, rather than sorting or translational inhibition strategy, or subtype identity.

All four libraries display additional quality control indicators of appropriate sequence-specificity. They each possess distinctly narrower length distributions over CDS compared to 3’ UTRs and snoRNAs (Figure 4E). Further, all four libraries exhibit a clear 3-nt P-site periodicity (Figure 4F), and strong enrichment for in-frame P-sites (Figure 4G). Interestingly, the SCPN sample displays higher P-site frequency closer to stop codons than to start codons, whereas the CPN libraries all display a slight preference for P-sites closer to start codons than to stop codons (Figure 4G). It is likely that this arises because the SCPN sample was sorted in the absence of translational inhibitor, and the distribution reflects elongation dynamics not present in the CPN samples treated with cycloheximide from tissue dissection onward.

Taken together, the data from these four independent samples from two distinct subtypes, with varying neuron number and [RNase I], and using two translation inhibition regimens, demonstrate rigorous reproducibility with as few as 17K sorted neurons. All samples meet the three critical quality control metrics (enrichment over CDS, distinct length distributions over CDS, and strong 3-nt periodicities) indicating that they capture translating ribosome footprints.

### nanoRibo-seq QC metrics remain robust using as few as 2.5K cells

We next further investigated the impact of cell number on Ribo-seq QC metrics, and aimed to identify a practical “lower bound” of input cell number for production of successful nanoRibo-seq libraries. We FACS-purified IUE-labeled CPN at P4 from individual labeled cortical hemispheres, performed nanoRibo-seq (RP, [RNase I]=12.5 U/mL) and alkaline fragmentation (AF), producing a range of 2.5-20.8K sorted neurons between samples (Figure 4-figure supplement 1C). All RP libraries display enrichment for CDS compared to 5’ and 3’ UTRs (Figure 5A), very consistent footprint length distributions (Figure 5B), and similarly distinct length distributions over CDS compared with 3’ UTR and snoRNA sequence (Figure 5C). These results indicate highly reproducible digestion patterns an over 8X range of cell numbers. In addition, all samples exhibit a strong 3-nt periodicity (Figure 5D), and have strong enrichment for in-frame P-sites (Figure 5E), strongly indicating that libraries are of high quality, and capture translating ribosomes with as few as 2.5k cells.

**Figure 5:**
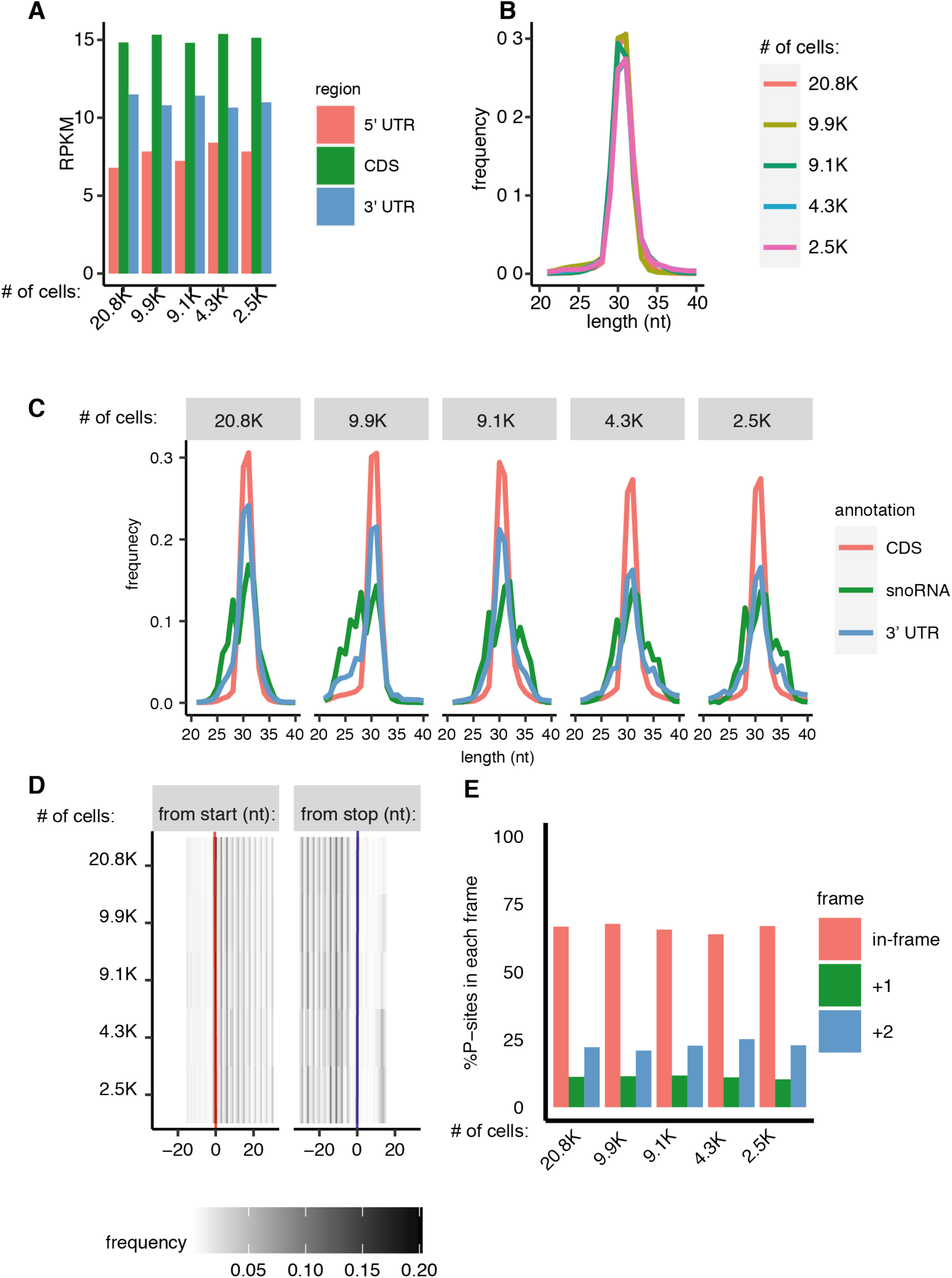
nanoRibo-seq from CPN purified out of individual labeled cortical hemispheres. A) RPKM coverage (reads/kilobase)/(million uniquely mapped) over CDS (5’ UTR=red, CDS=green, 3’ UTR=blue). B) Fragment length distributions over CDS. C) Fragment length distributions over CDS (red), snoRNAs (green), 3’ UTR (blue). D) Heatmap of P-site signal counts as a function of position from the start (red) or stop codon (blue). E) % of P-sites mapping to each reading frame (in-frame=red, +1=green, +2=blue).

### nanoRibo-seq measures RPF abundance for thousands of mRNAs, with high reproducibility between samples

An additional consideration for evaluating low input approaches to investigation of translational regulation is the ability to robustly detect and quantify translation of a large proportion of the transcriptome. To assess how many genes receive sufficient read depth for quantitative analysis, we examined the distributions of gene-level RPF abundances. Importantly, the QiA-seq miRNA kit used to generate the libraries incorporates a 12-nt random barcode at the 5’ end of Read 2, thus enabling identification and removal of PCR duplicates before performing all analyses. Gene-level raw count distributions reveal that raw counts span nearly four orders of magnitude in the highest input samples, decreasing to three orders of magnitude in the lower-input samples (Figure 6A). These distributions indicate that many thousands of genes receive 10s-1000s of reads, even in the lowest input cases.

**Figure 6:**
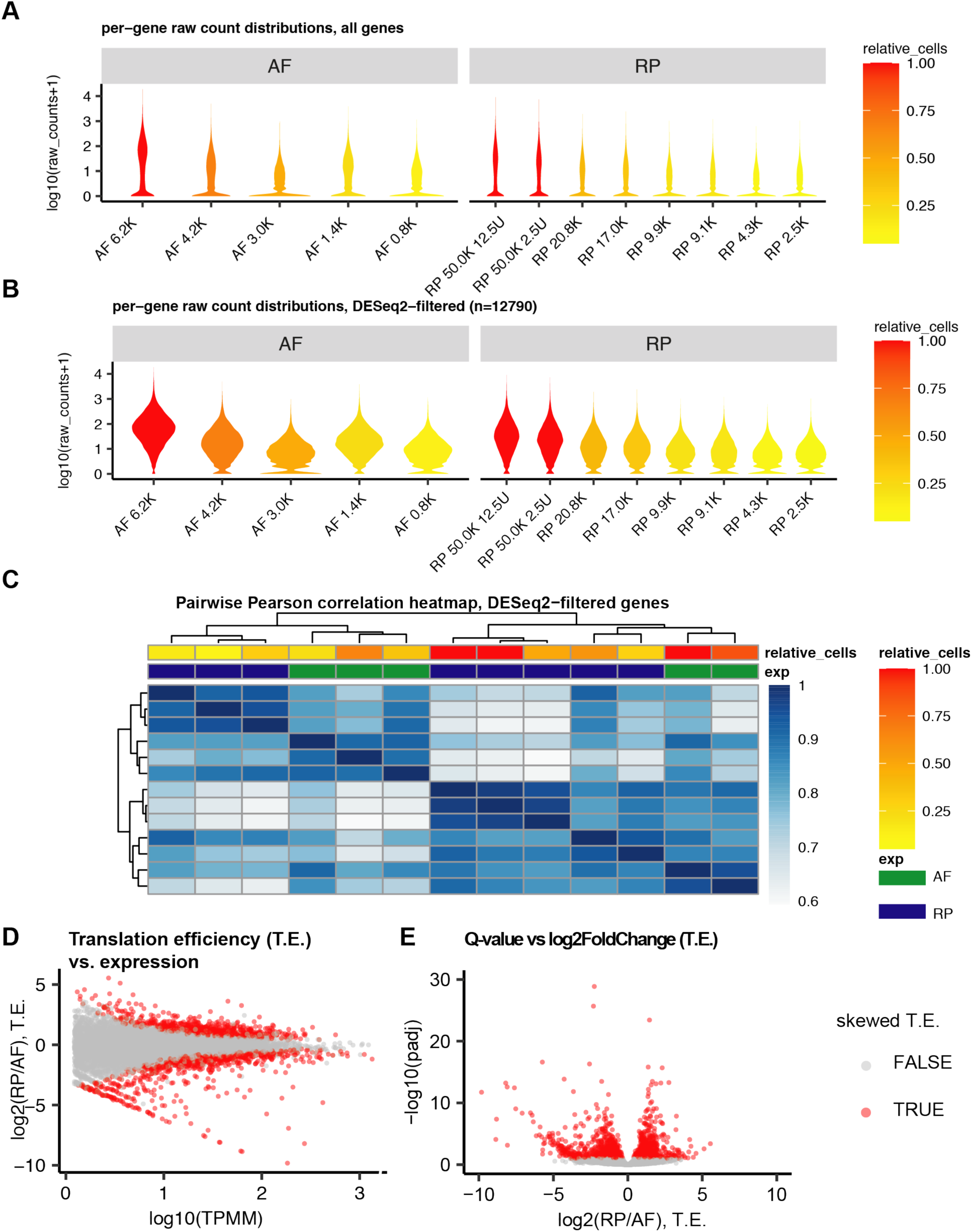
Detection sensitivity, sample clustering, calculation of translation efficiency, and identification of genes with skewed T.E. A) Per-gene raw count distributions from each sample, for all genes, colored by input cell quantities. Input cell quantities are depicted as “relative_cells”, the ratio of cells in a sample to the maximum number of cells used in the same experiment type (e.g., the max number of cells in the AF experiments is 6.2K, so the highest AF sample (6.2K cells) has relative_cells=1.0, and the lowest AF sample (0.8K cells) has relative_cells=0.13). Red corresponds to highest relative cell input quantities, orange to medium input, and yellow to the lowest input. B) Raw count distributions from each sample for the 12,790 genes passing DESeq2 independent filtering. C) Heatmap and hierarchical clustering based on the Pearson correlation coefficients of TPM expression values. The annotation columns depict relative_cells (red to yellow), and experiment type (green=AF, blue=RP). D) Translation efficiency (log_2_(RP/AF)) vs. expression (mean TPMM across all samples) for all genes passing DESeq2 filtering. Red indicates that a gene has a significantly skewed T.E. E) –log10(adjusted p-value) vs. T.E., (log_2_(RP/AF)). Red indicates that a gene has a significantly skewed T.E.

We next filtered out genes with expression too low to be informative for statistical comparisons between RP and mRNA abundance, by removing genes that failed to pass DESeq2’s independent filtering cutoffs. This left 12790 genes for analysis; distribution plots of the raw counts for genes passing the DESeq2 filter showed that, after the lowest expressed genes in each sample were removed, in all cases approximately half of the remaining genes in each sample are supported by at least 10 raw counts (Figure 6B). These results indicate that using the unbiased DESeq2 independent filtering approach both removes the least reliably expressed genes, and still leaves over 10,000 genes available for analysis, supported by robust numbers of raw reads even in the lowest input samples.

Another consideration is the reproducibility between experiments. To examine this, we calculated Pearson’s correlation coefficients, comparing depth and gene-length normalized counts (transcripts per million mapped, TPM) for all combinations of CPN libraries, using the 12790 genes above the DESeq2 independent filtering threshold. We then used these pairwise correlations as the distance metric for hierarchical clustering.

We find that RP libraries and AF libraries cluster separately from each other, as expected given the potentially large differences in transcriptional and translational output (Figure 6C). It is interesting to note that, despite their differences, two of the three lowest input RP libraries and the three lowest input AF libraries cluster separately from the higher input libraries (Figure 6C), which cluster together despite being generated from distinct subtypes isolated at slightly different developmental stages (P3 vs P4) after electroporation at slightly different ages (E14.5 vs E15.5). The 9.1K RP sample clusters with the higher input samples, while the 9.9K RP sample clusters with the lower 4.3K and 2.5K RP samples. This clustering pattern suggests substantial reproducibility between the higher input samples, despite distinct experimental batches and modest differences in the developmental timing of the samples. In contrast, there was less uniformity among the lowest input samples. Importantly, each data point in this analysis represents one unique biological sample, so there is potentially both technical variability due to varying numbers of cells, and biological differences between the samples.

Together, these data indicate that ∼10K and ∼3K cells are estimates for the input optimal for extremely highly reproducible RP and AF libraries, respectively. However, the data also indicate that, for biological systems limited to fewer than 10K cells, application of nanoRibo-seq with its strong Ribo-seq QC metrics likely would still provide informative translation efficiency estimates, especially for more highly abundant genes.

### Translation efficiency (T.E.) measurement reveals that approximately 10% of the CPN transcriptome has divergent RPF and mRNA abundance

To investigate translational regulation via calculation of T.E., in particular to assess rates of individually regulated protein translation independent of simple mRNA transcript abundance in CPN, we performed differential abundance analysis using DESeq2 between the five paired CPN RP and AF libraries generated from single hemispheres. This differential abundance analysis compares RPF abundance from the RP experiments to mRNA abundance in the AF experiments, thus both enabling calculation of translation efficiency T.E. as log_2_(RP/AF), and identifying the set of genes with significantly higher or lower than standard T.E. (i.e. divergence of the ratio of RPF abundance to mRNA abundance compared to most transcripts).

We find that 9.5% of all genes passing independent filtering display a significant discrepancy of the ration of RP abundance and mRNA abundance compared to most transcripts (Figure 6D & E). There are slightly more genes with a significantly low T.E. (T.E. < 0, n=873 genes) than genes with a significantly high T.E. (T.E. > 0, n=762 genes). Together, these results suggest that about one tenth of the CPN transcriptome displays significant divergence between RPF abundance and the respective mRNA abundance, indicating that a relatively large proportion of genes regulate protein product abundance via post-transcriptional, translational regulation in CPN.

### uORF translation by CPN: transcripts coding for synaptic and neuronal adhesion genes are enriched among mRNAs with translated uORFs

Upstream open reading frames (uORFs) are short open reading frames located in the 5’UTRs of approximately half of all mammalian mRNAs[25, 26, 39]. Though uORFs are often inefficiently translated because they have either poor translation initiation contexts[25] and/or use “near-cognate” non-canonical start codons[21], translation of a uORF has substantial consequences, since it is frequently inhibitory for translation of the main CDS[25, 26, 39] via a variety of mechanisms[37]. Since most uORFs are not constitutively translated[25, 26], changes in uORF translation might enable tight control of translation of specific transcripts in response to varying cellular conditions, in a cell type-specific manner. Thus, identifying which uORFs are translated by distinct cell types provides insight into which mRNAs might be especially subject to translational regulation by uORFs.

To identify all ORFs– novel or annotated– that display evidence of translation by CPN, we pooled read data (after duplicate removal) from all three pilot CPN replicates and the five single-hemisphere libraries, and used the RiboCode analysis package[40]. RiboCode identifies translated ORFs by testing whether there is a statistically significant bias for inferred P-sites in-frame with an individual ORF, compared to out of-frame P-sites[40]. This analysis identified 12390 translated, previously annotated protein-coding ORFs in the CPN data sets, and additionally identified 1698 uORFs, 517 “overlap uORFs” (Fig 7A), and over one thousand novel ORFs in other categories.

**Figure 7:**
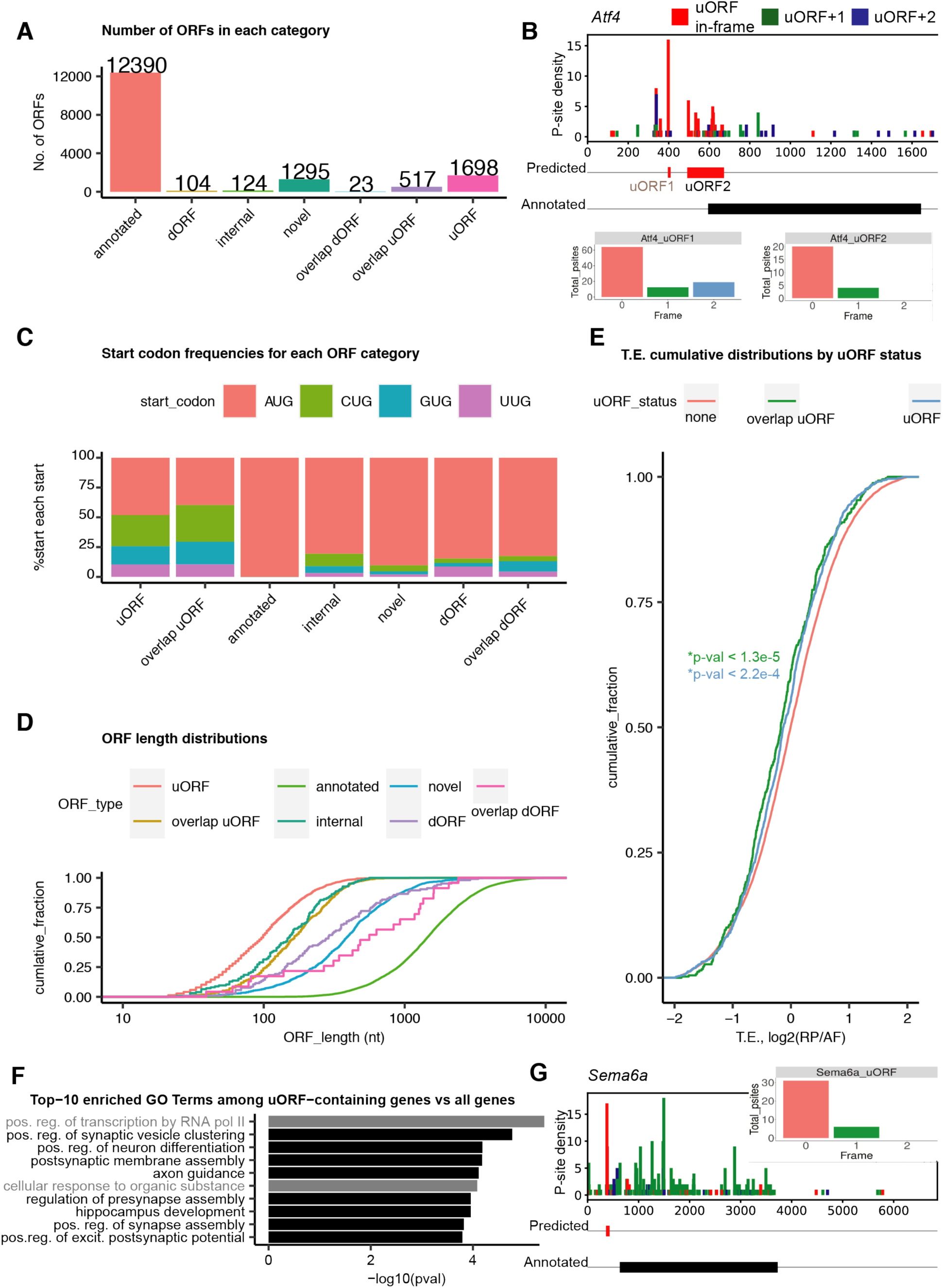
uORFs are enriched among mRNAs encoding proteins involved in synapse formation and neuronal adhesion. A) Numbers of each type of ORF identified as translated by RiboCode across all three CPN samples. B) P-site density as a function of position across the Atf4 mRNA transcript. P-site densities are color-coded by reading frame. The red box below the density plot shows the position of the predicted uORFs; the black box shows the annotated CDS (regions before and after are 5’ and 3’ UTRs, respectively). Inset boxes: the number of P-sites in reading frame: 0 (in-frame, red), +1 (green), +2 (blue) for the two Atf4 uORFs C) Start codon distributions by ORF category. D) Length distributions by ORF category. E) Cumulative distributions of T.E. for genes with no translated uORF (red), genes with overlap_uORFs (green), and genes with translated uORFs (blue). P-values were calculated using the Wilcox ranked sum test comparing T.E.s between genes with uORFs (green) or overlap_uORFs (blue) with genes lacking called translated uORFs. F) –log10(p-values) for top-10 enriched biological process GO terms among uORF- and overlap uORF-containing genes; GO terms related to synaptic biology and/or neuronal circuit formation are highlighted in red. G) P-site density and sum in each frame for the Sema6a uORF. Inset boxes: the number of P-sites in reading frame: 0 (in-frame, red), +1 (green), +2 (blue) for the Sema6a uORF.

For example, these data independently highlight translation of two already known, highly conserved uORFs in the 5’ UTR of Atf4. Atf4 uORF1 is a short ORF, just 3 amino acids long, which is translated under non-stressed conditions, and which promotes translation of uORF2. Atf4 uORF2 is a longer “overlap uORF”, which is highly inhibitory to Atf4 CDS translation under non-stressed conditions[41]. RiboCode analysis identifies uORF2 as a uORF that is translated by CPN, with strong evidence of enrichment for in-frame P-sites (Figure 7B). Though uORF1 is too short to be called by RiboCode (we used a >6 AA length filter), it too displays strong enrichment for in-frame P-sites (Figure 7B), suggesting that uORF1 is also translated by CPN. These results demonstrate the ability of nanoRibo-seq to identify uORF translation– e.g., detecting well-established individual examples of Atf4 uORF translation, thus providing foundation and validation for identification and analysis of other uORFs and novel ORFs.

Importantly, the overall number, length, and start codon characteristics of uORFs identified as being translated in these analyses are generally consistent with measures of these parameters in translated uORFs identified in other vertebrate Ribo-seq analyses[21, 42]. In particular, approximately half of uORFs and “overlap uORFs” (uORFs that partially overlap start codons) identified here use near-cognate “NTG” start codons (Figure 7C), and the translated uORFs and overlap uORFs identified are short, on average ∼100- and 200-nt long, respectively (Figure 7D). Interestingly, compared to genes without translated uORFs (mean T.E.=1.15), genes containing either translated uORFs or overlap_uORFs exhibited on average decreased T.E.s (those with translated uORFs: mean T.E.=1.03, p-value < 1.3e-5; those with translated overlap uORFs: mean T.E.=1.02, p-value < 2.2e-4) (Figure 7E). These results are consistent with the typically inhibitory role of uORF translation on CDS translation in CPN.

We next investigated key biological characteristics of genes containing uORFs. We first performed a gene ontology (GO) analysis, using genes with uORFs or “overlap-uORFs” translated by CPN as the gene list, and genes with annotated protein coding ORFs translated by CPN as the background. Intriguingly, we found strong enrichment of translated uORF-containing transcripts (against the background of all translated, annotated protein coding transcripts expressed by CPN) with GO terms for biological processes including pre- and post-synaptic membrane assembly, axon guidance, and neuronal development (Figure 7F). This suggests that uORFs in genes involved in neuronal synapse and circuit formation are active during early postnatal CPN circuit development, and might function to refine regulation of, and potentially more tightly control, synthesis of important functional proteins from these synaptic transcripts.

There are many examples of translated uORFs in transcripts involved in biological processes highly relevant for CPN identity, and with functional roles in axon guidance, synapse formation, and neuronal adhesion needed to form proper cortical circuitry. mRNAs encoding TFs are enriched among uORF-containing genes (Figure 7F), we identify dozens of TFs with translated uORFs, including TFs previously known to be critical for the specification of superficial layer CPN identity and circuitry, e.g. Cux1[43], Bcl11a[44, 45], Satb2 [46, 47], and Cited2[48] (Figure 7-figure supplement 1). We also identify a translated uORF in Sema6a mRNA (Figure 7G), a member of the semaphorin class of receptors, which plays key roles in axon guidance[49]. Intriguingly, we find that genes for the neuronal adhesion and synapse selection proteins neuroligin 1,2, and 3 all have multiple translated uORFs (Figure 7-figure supplement 2).

Extending this set of functional clusters involved in circuit development, mRNAs coding for neurexin protein family members display a distinct pattern of uORF translation. Neurexins are synaptic membrane proteins with a wide set of interaction partners, including neuroligins, with crucial roles in synapse selection, establishment, and maintenance[50]. Neurexins are encoded by three genes (Nrxn1,2,3), each with α- β- and γ-isoforms differing by alternate transcription start sites and 5’ UTRs. Further, each gene contains multiple alternatively spliced exons[51], leading to a complex “molecular code” that favors binding interactions between particular neurexin isoforms and specific binding partners at particular synapses [50]. Nrxn1α, Nrxn1β, and Nrxn3α 5’ UTRs contain multiple translated uORFs (Figure 7-figure supplement 3) in developing CPN (Nrxn1α: 6 translated uORFs, one of which occurs in a distinct 5’ UTR from an alternate transcription start, and Nrxn1β: 3 translated uORFs). Intriguingly, the Nrxn1α 5’ UTR harbors more translated uORFs than any other 5’UTR identified in this analysis. Previous work found that multiple conserved, short uORFs within the Nrxn2α 5’UTR cooperatively inhibit translation *in vitro* (Figure 7-figure supplement 4A)[52]. Interestingly, we find that a longer Nrxn2α uORF using a non-canonical CUG start codon is also translated in developing CPN (Figure 7-figure supplement 4A). While the uORFs identified in the previous study[52] are too short to be directly called by RiboCode, plotting P-site density across these uORFs reveals a bias toward in-frame P-sites across all three uORFs, especially uORF2.

These results suggest that these uORFs are likely also translated by CPN (Figure 7-figure supplement 4A), extending previous findings to an *in vivo* setting. In mouse, the CUG uORF would be in-frame with uORF2, but is interrupted by a stop codon (Figure 7-figure supplement 4B). In humans, there is also a longer CUG uORF that reads through into uORF2, and actually encompasses the entire uORF2 sequence (Figure 7-figure supplement 4C). The peptide sequence of the human CUG uORF contains a region of moderate similarity to the mouse CUG uORF, and both are rich in Gly and Pro residues, both containing 13 Pro residues (Figure 7-figure supplement 4D). These findings suggest possible conservation of this CUG uORF, and similar polyproline peptide sequences imply that it might induce ribosome stalling/pausing[53, 54].

The identification of multiple uORFs in Nrxn1α, β, Nrxn2α, and Nrxn3α 5’ UTRs, along with previous molecular dissection of short uORFs with canonical start codons in Nrxn2α[52], suggests that uORFs might be translational regulators neurexin. Taken together, our results identifying uORF translation by developing CPN suggest that, during early postnatal CPN development and circuit formation, many genes with translated uORFs function as regulators of axon guidance, cell adhesion, and synapse selection/formation, including multiple examples of semaphorins, neuroligins, and neurexin.

## DISCUSSION

nanoRibo-seq is a highly effective, quantitative, optimized approach for ultra-low-input ribosomal profiling (Ribo-seq) that requires 10^2-^10^3^ less input material than current approaches for comprehensive investigation of sets of mRNAs undergoing active translation. The resulting ribosomal profiles display highly stringent quality control across a range of cellular input from as few as 2.5K cells. We present nanoRibo-seq results and their reliable quality control metrics from a range of cellular input and RNaseI digestion conditions, using FACS-purified, subtype-specific cerebral cortex (long-distance circuit) “projection neurons” as a model system.

We first demonstrate the feasibility and optimized conditions for nanoRibo-seq with as little as 30 ng RNA brain lysate material, by testing whether the global read distributions obtained conform to three expected features of successful Ribo-seq data sets: 1) enrichment of CDS compared to UTRs and non-coding RNA; 2) distinct length distributions over CDS and ncRNA; and 3) strong 3-nt, in-frame inferred P-site periodicity. All three quality control metrics are stringently and reproducibly achieved. We also present several approaches for generating randomly fragmented control libraries, favoring alkaline fragmentation for its simplicity and tight size distributions.

We then investigated whether nanoRibo-seq could produce high-quality translational profiles from small, experimentally limited quantities of FACS-purified, subtype-specific callosal projection neurons (CPN), and subcerebral projection neurons (SCPN). We obtained high nanoRibo-seq quality libraries from just 2.5K neurons, with quality assessed both in terms of the expected sequence features of Ribo-seq, and in terms of robust and comprehensive quantification of RPF abundance for thousands of genes. Though our data suggest that using at least 10K neurons is optimal for reproducibility across samples, high quality and informative data result from as few as 2.5K neurons.

We employ nanoRibo-seq to generate transcript-specific translational efficiencies (T.E.s) to investigate potential transcript-specific translational regulation. We compare RPF abundance to mRNA abundance across all transcripts, and find that ∼10% of detected genes display statistically significant divergence of RPF abundance to mRNA abundance ratios compared with the total transcript population, suggesting that this substantial ∼10% portion of the CPN transcriptome undergoes transcript-specific positive or negative translational regulation.

Intriguingly, nanoRibo-seq data from CPN *in vivo* identify a large number of actively translated uORFs. The translated uORF data identify enrichment for genes involved in key processes for neuronal circuit formation and maturation; neuronal adhesion, pre- and post-synaptic membrane assembly, and synaptic transmission are enriched among transcripts containing translated uORFs. Multiple examples of translated uORFs are present in the 5’ UTRs of key regulators of synapse selection, maintenance, and activity, including semaphorins, neuroligins, and neurexins. The uORFs within neurexins provide important grounding in known biology from *in vitro* studies, and extend importantly from what was previously known– our data indicate *in vivo* translation of multiple uORFs within Nrxn2α previously shown to be potent regulators of its translation *in vitro*[62], along with discovery of *in vivo* translation of a novel CUG-start uORF coding for a conserved, proline-rich peptide. Further, we identify that Nrxn1α and Nrxn1β 5’ UTRs also harbor a complex set of multiple translated uORFs, indicating that uORF translation by CPN occurs more broadly, within multiple neurexin isoforms. These and other translated uORFs in the 5’ UTRs of multiple key regulators of development and maintenance of neuronal circuitry and synaptic connectivity suggest refinement and potentially tightly regulated translation of these mRNAs by developing CPN via uORF translational activity. It would seem likely that related mechanisms might be employed widely by many neuron subtypes, and potentially much more widely by other types and subtypes of cells across tissues and organ systems.

We highlight uORFs as examples of non-canonical translation uncovered using nanoRibo-seq because of their well-studied roles in translational regulation of downstream coding ORFs, though we note that this is only one possible role for uORF translation. We also identified over a thousand examples of other non-canonical translated ORFs, and both these novel ORFs and uORFs might produce a variety of translation products with novel biological functions. Interestingly, and potentially directly related to our results, a recent RiboSeq analysis of human neocortex also identified a wide array of translated small ORFs, some of which could be validated as encoding micropeptides by mass spectrometry, and many of which are primate- or human-specific[32]. These observations in humans, combined with our results from mice reported here, suggest a large set of novel translated ORFs in the mammalian cortex. nanoRibo-seq will likely be a powerful and extremely informative approach for further interrogation of these novel translation events by specific neuronal subtypes, likely at specific developmental stages, at specific neuronal activity and environmental states, and under other specific biological contexts.

nanoRibo-seq enables investigation of translational regulation in a wide variety of previously inaccessible biological contexts. Standard Ribo-seq approaches require large amounts of input material. nanoRibo-seq requires only a few thousand cells, making possible ribosomal profiling and translational regulation investigation in highly specific and/or rare populations of cells. It is also a complementary approach to a recently-developed single-cell Ribo-seq method[36], in that it enables measurements of T.E.s in populations of interest (as opposed to simply measuring RPF abundance without normalizing to mRNA abundance). Importantly, and offering broad applicability and ease, nanoRibo-seq requires no specialized equipment, robotics, or microfluidics.

nanoRibo-seq offers benefits compared with common techniques that examine translational output from specific cell types by identifying ribosome-bound transcripts through Cre-dependent expression of tagged ribosomal proteins[55–57]. These approaches rely on specific driver lines, available only in some model systems. nanoRibo-seq does not require genetic labels, and is thus amenable to any labeling strategy compatible with FACS sorting or other cellular enrichment or purification methods. Further, as demonstrated here by the identification and analysis of extensive uORF translation by CPN in the 5’UTRs of key synaptic structural and regulatory genes, nanoRibo-seq enables single nucleotide-level analysis and discovery of novel patterns of translation and/or novel translational regulatory mechanisms. Importantly for practical and widespread application, a broad range of digestion conditions yield high-quality Ribo-seq and control alkaline fragmented libraries, making generalization to other cell types and tissues straightforward. The results presented here predict that nanoRibo-seq will be extremely useful for investigation of translation and translational regulation by many specific cell types during their development and maturation, in disease models, in non-model organisms lacking a broad set of genetic tools, in specific subcellular compartments of neurons and other cell types, and from small human patient samples.

## Contributions

J.E.F and J.D.M conceived of experiments, and wrote the manuscript. JF performed all Ribo-seq experiments, and computational analysis. O,D performed IUE labeling.

## Acknowledgements

We thank Dr. Yasuhiro Itoh and Melody Ross for a careful reading and discussion of this manuscript. This work was supported by NIH DP1 NS106665, with additional infrastructure support from R01 NS104055, R01 NS045523, and the Max and Anne Wien Professor of Life Sciences fund. J.F. was partially supported by T32 AG000222, F32 AG067661, and the DEARS Foundation (to JDM).

## Competing Interests

None.

## Data and Code Accessibility

Sequence data has been deposited to GEO Accession GSE197060. Code is available on Github at at https://github.com/6LayeredCortex/nanoRibo-Seq

## METHODS

### Mice

The vertebrate animal experimental protocols were approved by the Harvard University Institutional Animal Care and Use Committee. For *in utero* electroporation experiments, we ordered timed pregnant CD-1 dams from Charles River Laboratories, and performed *in utero* electroporation of fluorescent labeling constructs at E14.5 or E15.5. For SCPN retrograde labeling experiments, we bred B6SJL x CD-1 mice, and performed ultrasound-guided injections into the cerebral peduncle at P1.

### Neuronal Labeling

#### CPN

We performed electroporation of fluorescent constructs using CD-1 mice at E14.5 or E15.5. Briefly, we exposed both uterine horns, and injected a solution of 5 ug/uL DNA + 0.1% FastGreen in 1X TE containing 5ug/uL plasmid encoding myristolated-TdTomato into one lateral ventricle in approximately one half of embryos. We then electroporated each injected embryo with 5 pulses of 500 ms each at 34 mV. We screened P1 pups for red fluorescence in the cortex under an epifluorescence dissection microscope.

#### SCPN

We labeled SCPN by ultrasound-guided injection into the cerebral peduncles at P1 of B6SJL x CD-1 pups bred in our colony, using injection conditions described previously[45, 58]. Briefly, we anaesthetized the pups hypothermically by placing them on ice for 3 minutes, then gently stabilizing them on the injection platform. We visualized the cerebral peduncle and placement of the injection micropipette via ultrasound backscatter microscopy. We injected Alexa 555-cholera toxin B (CTB) at four sites in the left cerebral peduncle (dorsal, ventral, medial, and lateral sites) to ensure coverage of the peduncle, with five 5 nl pulses per site.

### Neuronal FACS

We performed neuronal FACS using established approaches[45, 58–62], but with the addition of translational inhibitors to prevent *ex vivo* translational elongation; for the CPN sorts, all buffers contained 100 ug/mL cycloheximide (CHX). We dissected brains into HBSS buffer + CHX, visualized the labeled region of cortex using an epifluorescence dissection microscope, and dissected the labeled cortical region into disociation solution (DS) + CHX. We washed cortical tissue pieces twice in DS + CHX, then enzymatically digested twice by incubation in enzyme solution (DS with cysteine and papain) + CHX for 15 minutes each time (with inversion every 5 minutes to keep tissue pieces mixed). We washed twice in wash solution (WS) + CHX, triturated 15-20 times in ∼1 mL of WS + CHX using fire-polished glass pipettes. We diluted the cell suspension with 4 mL WS + CHX, spun down cells for 5 minutes at 1000g, triturated again in 1 mL WS+CHX, and passed the cell suspension through a strainer cap. We added 1:1000 SYTOX Blue to enable screening out dead cells. We then FAC-sorted the red cells, and collected them in a DNA LoBind Eppendorf tube containing 2X polysome buffer + CHX (25 mM HEPES pH=8.0, 200 mM KCl, 10 mM MgCl_2_, 4 mM CaCl_2_, 1% nonidet P-40, 100 ug/mL CHX). For the SCPN sort, we performed all steps as described above, except that we omitted CHX until the SCPN were sorted into 2X polysome buffer + CHX.

### Generation of Ribosome Protected Fragments (RPFs), general approach

To perform RNase I digestion, we diluted brain lysate or FACS-purified neurons into 1X polysome digestion buffer (2X polysome buffer diluted 1:1 with dilution buffer, which is 0.75X PBS, 0.08M sucrose, 100 ug/mL CHX, 1/500X RNasin), added the specific amount of RNase I for each experimental variation (Ambion brand, ThermoFisher #2294), and placed the mixture on a rocker for 45 minutes at room temperature. To stop digestion, and concentrate the very dilute RNA for purification, we added 10 uL Superasin (200 units, Ambion brand, ThermoFisher #2694), poured the reaction into an Amicon Ultra-15 Centrifugal Filter (10kD NMWL filter, EMD Millipore UFC901024), and centrifuged at 5000g at 4°C for approximately 30-60 minutes, until <400 uL of retenate remained. To purify the RNA, we used a slight modification of the Zymo Quick RNA Microprep (Zymo Research #1051) extraction kit procedure to purify small RNAs out of large volumes. We transferred the retained RNA to a 15 mL Falcon tube, added 800 uL lysis buffer (mixing vigorously by pipette), then added 4 mL 100% EtOH (mixing by pipette or inversion at least 7 times). We then repeatedly centrifuged the solution onto a single Zymo spin column (Zymo Spin IC Column, Zymo Research #C1004), spinning at full speed for 15 seconds in a microcentrifuge until all solution from one sample had been passed over that single column. We then followed the remainder of the Zymo RNA preparation protocol; in brief: we spun the column with 400 uL prep buffer, then 700 uL wash buffer, 400 uL wash buffer, and spun the column empty to remove residual wash buffer. We then transferred the column to a labeled DNA LoBind Eppendorf 1.5 mL tube (VWR # 022431021), added 20 uL nuclease-free H_2_O, allowed the column to sit for 5-10 minutes at room temperature, then spun at full speed on a microcentrifuge. The eluted RNA is able to be frozen at -80 for long-term storage.

To enrich RNAs in the RPF size range, we gel purified the RNA fragments. First, we added 20 uL 2X formamide denaturing solution (20% Glycerol, 80% Formamide (Promega # H5051), 60 mg/mL Bromophenol Blue) to the RNAs, heated at 70°C for 5 minutes, placed on ice until ready, then loaded onto a 15% TBE-UREA acrylamide gel, and electrophoresed in 1X TBE running buffer at 200V for approximately 45 minutes (until the bromophenol blue marker reached the bottom of the gel). To ensure proper sizing, we ran 5 ng 10/60 Oligo Length Standards 10/60 Ladder (IDT # 51-05-15-01) and 1-3 ng of synthetic 27 and 36 nt RNA oligos. We stained the RNA in gels with SYBRGold (Thermo S11494) for 10 minutes in 1X TBE, followed by a 5 minute wash with 1X TBE, and two rinses with 1X TBE to remove background. We then imaged RNA in the gels using a standard gel documentation instrument (BioDoc-IT Imaging System, Analytik Jena) with UV transillumination, exposing for 20 seconds. We excised the RNA running between the 27-36 nt markers. To purify the RNA, we placed the gel slice in 400 uL Crush and Soak buffer (300 mM KOAc, 1 mM EDTA, 1% SDS) in a LoBind Eppendorf Tube, and rotated on a shaker at room temperature overnight. We transferred the liquid to a 15 mL Falcon tube, added 400 uL Zymo Lysis Buffer and 4 mL 100% EtOH, repeatedly centrifuged one sample onto one spin column, then purified RNA using the standard Zymo kit protocol from the 400 uL prep buffer step onwards after binding the RNA to the column as described above.

Since the digestion conditions varied between experiments, some specific conditions varied across experiments:

#### 30 ng lysate initial experiment

To prepare the homogenate for the 30 ng initial trial, as well as for the bulk trial, we harvested three P4 CD-1 forebrains (olfactory bulbs removed; midbrain, cerebellum, and brainstem removed via vertical cut at caudal end of cortex) into ice-cold 1X polysome buffer+CHX, homogenized in a dounce homogenizer with 11 strokes on ice, transferred to a 15 mL Falcon tube, and centrifuged at 1660g at 4°C for 15 minutes to pellet nuclei. We then extracted RNA from 60 uL of homogenate using the standard Zymo Quick RNA protocol, and measured RNA concentration via a Nanodrop (Nanodrop 1000, Thermo). From this estimate of RNA concentration, we calculated the volume of homogenate equivalent to 30 ng of RNA. We added this volume of homogenate to 6 mL of 1X polysome digestion buffer, added RNase I to a final concentration of 2.5 U/mL, and allowed digestion to proceed for 45 minutes on a rocking platform covered in aluminum foil to prevent photodegradation of cycloheximide. We then added 10 uL Superasin, and purified footprints as described above.

#### Bulk experiment

We used the same lysate as used for the 30 ng expriment, and digested ∼120 ug RNA equivalent (640 uL) with 1500 U RNase I in 640 uL for 45 minutes at RT on a rocker covered in foil. We added Superasin, then split the lysate across 6 separate columns, purified RNA using the standard Zymo Quick RNA Microprep protocol, and combined elutions into one tube. We measured the RNA concentration of this bulk footprint elution using a Nanodrop, and ran the equivalent volume of 1 ug on PAGE gel to purify footprints.

#### RNase I titration

We homogenized P6 forebrains in 10X volume of 1X polysome buffer+CHX, as described in the 30 ng and bulk initial trials. We extracted RNA from 60 uL homogenate, measured the RNA concentration, and calculated the volume of lysate equivalent to 30 ng RNA. We added this volume to four separate reactions consisting of 6 mL 1X polysome digestion buffer, and either 2.5, 12.5, 62.5 or 312.5 U/mL RNase I. We allowed digestions to proceed for 45 minutes at room temperature on a rocking platform covered in foil, added Superasin, and extracted footprints as described above.

#### sorted SCPN

We labeled SCPN via injection of Alexa-555 conjugated CTB into the cerebral peduncle of B6SJL x CD-1 P1 mice as described above. We harvested P3 brains, then dissected and dissociated cortices for neuronal FACS using standard FACS without cycloheximide added during these steps. We sorted Alexa-555-positive neurons into 2X polysome buffer+CHX, with added RNaseIn. We diluted the volume equivalent of 50,000 sorted neurons into a final volume of 4.1 mL 1X polysome digestion buffer+CHX, and digested with 12.5 U/mL RNase I for 45 minutes at room temperature on a foil-covered rocking platform. We added Superasin, and extracted footprints as described above.

#### sorted CPN

We labeled CPN via *in utero* electroporation of myristoylated-TdTomato plasmids into E14.5 embryos. We harvested P3 brains into ice-cold HBSS + 100 ug/mL cyclohexmide, dissected the labeled cortical hemisphere, and dissociated it for neuronal FACS using our established protocol except with 100 ug/mL cycloheximide added to all solutions to keep translational elongation inhibited. We sorted red fluorescent neurons into 2X polysome digestion buffer+CHX, diluted either 50,000 or 17,000 neurons into 4.1 mL 1X polysome digestion buffer+CHX, and digested with 2.5 or 12.5 U/mL RNase I for 45 minutes at room temperature on a foil-covered rocking platform. We added Superasin, and extracted footprints as described above.

#### sorted CPN from individual cortical hemispheres

We labeled CPN via *in utero* electroporation of myristoylated-TdTomato plasmids into E15.5 embryos. We dissected P4 brains into ice-cold HBSS + 100 ug/mL cyclohexmide, keeping each individual brain as a separate sample, dissected each labeled cortical hemisphere, and dissociated it for neuronal FACS using our established protocol except with 100 ug/mL cycloheximide added to all solutions to keep translational elongation inhibited. We sorted red fluorescent neurons into 2X polysome digestion buffer+CHX, and diluted the sorted cells to a concentration of 12,500 cells/mL in 1X polysome digestion buffer following sorting. We set aside 1/5 of the sorted cells from each sample for RNA extraction, and preparation of the alkaline fragmented control libraries described below. We digested all samples with 12.5 U/mL RNase I for 45 minutes at room temperature on a foil-covered rocking platform. We then added Superasin, and extracted footprints as described above.

### Fragmentation of total RNA to produce control libraries

#### Rnase I digestion

##### G -> R method

We digested 30 ng purified forebrain RNA in 60 uL 1X polysome buffer with 1 uL RNaseIn and 0.16 U RNase I for 45 minutes at room temperature, then purified the RNA by first adding 90 uL lysis buffer+650 uL 100% EtOH, then following the Zymo Quick RNA Microprep protocol. We then gel purified the fragments between 20-60 nt as described for the ribosome footprints. We then carried out rRNA depletion with the NEBNext rRNA Depletion Kit (Human/Mouse/Rat NEB # E6310S) using the manufacturer’s instructions, however, we purified RNA at the end of the procedure using a Zymo Quick RNA Microprep cleanup (adding 90 uL Lysis buffer+650 uL 100% EtOH to the final reaction, then following the standard protocol thereafter) to purify the small RNAs at the last step of the NEB protocol instead of the suggested bead-based RNA cleanup protocol listed in the protocol.

##### R -> G method

We digested and purified 30 ng forebrain lysate RNA as described in the “G -> R” method. After digestion, we then carried out rRNA depletion with the NEBNext rRNA Depletion Kit as described above. We ran the purified RNAs after rRNA depletion out on a 15% TBE-Urea PAGE gel, and purified fragments from 27- to 60-nt using the previously-described gel purification procedure used to isolate ribosome footprints.

#### Alkaline Fragmentation

We performed the alkaline fragmentation time titration experiments using RNA extracted from whole dissociated P3 cortex from B6SJL x CD-1 mice. For one reaction, we mixed 7.5 ng RNA in 10 uL nuclease-free water with 10 uL bicarbonate fragmentation buffer (12 mM Na_2_CO_3_, 88 mM NaHCO_3_, pH=9.2) in a PCR strip, heated at 95°C for either 0, 5, 10, 15, 20, 25, 30, 35, or 40 minutes, and stopped the reaction by placing on ice, and adding 20 uL 2X formamide denaturing solution, and directly loading onto a 15% TBE-UREA PAGE gel immediately. We found that timepoints between 35-40 minutes produce the size distribution with the most RNA between ∼20- to 60-nt, in line with the input size range for the QiA-Seq miRNA Library Kit. We repeated the fragmentation for 37 minutes using 7.5 ng P3 cortex RNA, then gel purified fragments between ∼27- to 60-nt for library preparation.

To generate the alkaline fragmented libraries paired with Ribo-seq libraries from single cortical hemispheres, we set aside 1/5 of the material after sorting, extracted RNA using the Zymo Quick RNA Microprep protocol, and fragmented the eluted RNA by adding one volume of bicarbonate fragmentation buffer, heating for 37 minutes at 95°C, then gel purifying the fragmented RNA as described above.

### Oligo Length Standard Sequences

27 nt standard: AGUCACUUAGCGAUGUACACUGACUGA/3Phos/

36 nt standard: AUGUACACGGAGUCGAGCACCCGCAACGCGACUGUA/3Phos/

### Library Preparation

To ensure proper 5’P and 3’OH end moieties on the RNA fragments, we treated the RNA with polynucleotide kinase (PNK). We added 5 uL 1X T4 Ligase Buffer (NEB # B0202S), 0.5 uL T4 PNK (NEB #M0201S), 0.2 uL RNaseIn and 25 uL Nuclease-Free H_2_O to RNA samples, and incubated at 37°C for 30 minutes. We then added 80 uL Zymo Lysis Buffer, 650 uL 100% EtOH, and purified the RNA using the standard Zymo Quick RNA Microprep protocol, eluting in 14 uL. To estimate [RNA] and confirm purification of small RNAs, we ran the RNA on an RNA Pico Bioanalyzer Chip. We then used a SpeedVac to remove excess volume until RNA samples were under 5 uL. We prepared sequencing libraries using the QiA-seq miRNA Library Kit protocol as described, and we used the following dilutions of adapters and RT primers: 3’ adapter was diluted 1:5, 5’ adapter was diluted 1:2.5, RT primer was diluted 1:5. We amplified all libraries for 21 PCR cycles, and estimated library concentrations and size distributions with either the High-Sensitivity DNA or DNA Tapestation kits (we initially used the High-Sensitivity kit, but found that the library concentrations were often above the linear range of that kit).

### Sequencing

We pooled samples based on the library concentration estimates from the Tapestation runs. The Harvard Bauer Core performed final library quantification using KAPA-qPCR, and sequenced the experiments using the following machines and run setups: for the 30 ng initial lysate trial, bulk, RNase I in vitro digestion libraries: NextSeq Mid 2×75 bp reads; for the alkaline fragmentation, sorted CPN, sorted SCPN libraries, as well as the paired single-hemisphere CPN Ribo-seq and alkaline fragmentation libraries: NovaSeq SP 2×50 reads.

### Bioinformatic Analysis, and Ribo-seq Quality Control

#### Code and Data Accessibility

We have created a Github accession (https://github.com/6LayeredCortex/nanoRibo-Seq) containing all custom scripts used for this study. We performed all read processing, duplicate removal, rRNA and mm10 genome and transcriptomes alignment, length distributions, raw counting, RiboCode translated ORF identification using a Snakemake pipeline run as a batch job using Harvard University’s Cannon cluster. We performed RiboWaltz P-site periodicity analysis[38], TPM calculations, sample clustering, comparison, and sensitivity analysis, as well as uORF TopGo analysis using an Rmarkdown pipeline also run as a batch job (and as the final step in the Snakemake pipeline) on Harvard’s Cannon cluster. We include these Snakemake and Rmarkdown files, all needed inputs such as GTF files and Fasta files, and all custom UNIX shell scripts in the Github accession as full documentation of the computational methods used in this study.

#### Overview

Computational analyses were developed and performed to evaluate the following three Ribo-seq quality control metrics: 1) enrichment for CDS; 2) distinct length distributions over CDS compared to non-coding RNA; 3) enrichment for P-sites in-frame with annotated ORFs (Figure 1-figure supplement 1), as well as to 4) calculate translation efficiency; 5) identify genes displaying significantly skewed T.E. (Figure 6); and 6) investigate a potentially broad set of both annotated and novel ORFs (Figure 7). Importantly, we leveraged the low input library preparation approach using the QiA-seq miRNA kit to identify and remove PCR duplicates prior to analysis.

To accomplish these goals, we began by extracting the random barcode, which comprises the first 12-nt of Read 2 using umi-tools. We then aligned reads to rRNA using STAR, and filtered out the rRNA-aligned reads, as well as reads too short for unique alignment (<21-nt). We aligned the remainder to the mm10 genome and transcriptome with STAR. We next calculated CDS enrichment, and length distributions over CDS and other non-coding RNA, using custom shell scripts. We analyzed P-site periodicity and P-site frame enrichment with the R-package RiboWaltz[38]. We used RiboCode[40] to identify all translated ORFs, including novel, unannotated ORFs, such as uORFs[40], and we performed a gene ontology analysis to find gene sets enriched among uORF containing genes using the R-package TopGO. We calculated T.E.s by summing up read counts over CDS in RP data sets, and over entire genes for AF data sets for each gene using subRead:featureCounts, then performed filtering, normalization, and analysis of differential expression (between AF and RP data sets) with DESeq2.

#### Preprocessing, alignment, duplicate removal, and raw counting

To enable duplicate removal from these variable-length Ribo-seq sequences, we took advantage of the fact that we used paired-end sequencing, and that the first 12-nt of Read 2 of QiA-seq miRNA libraries is a 12-nt random barcode. We filtered for all reads with Read 2 > 12 reads (the vast majority), and extracted this random barcode using umi-tools: `umi_tools extract --extract-method=string --bc- pattern=NNNNNNNNNNNN`.

To remove the QiASeq miRNA adapters from the 3’ ends of Read 1, we ran Cutadapt v2.8.: `cutadapt -a AACTGTAGGCACCATCAAT`

We aligned Read 1 to the reference mouse rRNA sequences using STAR v2.6: STAR --runMode alignReads --runThreadN 8 --outSAMtype BAM Unsorted --readFilesCommand gunzip -c --outReadsUnmapped Fastx -- outSAMmultNmax 1 --outFilterScoreMinOverLread 0 -- outFilterMatchNminOverLread 0 --outFilterMatchNmin 21.

We then used the *final.out files produced by STAR for accounting the number and fraction of reads aligned to rRNA. We aligned the reads that failed to align to rRNA to the mm10 genome, and to the mm10 transcriptome for analysis of footprint length distribution, coverage over CDS, and producing gene-level raw counts. We aligned to the mm10 genome (GRCm38 build) using STAR v2.6 and the following parameters: STAR --sjdbGTFfile Mus_musculus.GRCm38.95_chrNamed_headFix.gtf -- runMode alignReads --runThreadN 8 --outSAMtype BAM SortedByCoordinate --quantMode TranscriptomeSAM -- readFilesCommand gunzip -c --outSAMmultNmax 1 -- outFilterScoreMinOverLread 0 --outFilterMatchNminOverLread 0 -- outFilterMatchNmin 21).

We completed duplicate removal following mm10 alignment to collapse all reads with the same alignment coordinates and exactly matching barcode sequences using umi-tools: `umi_tools dedup --method=unique`

We used the duplicate-removed bamFiles produced by this alignment for visualization, and for all further analysis. We then obtained raw, gene level counts over UTRs, CDS, and non-CDS using subRead:featureCounts, according to the annotations in Mus_musculus.GRCm38.95.gtf. We calculated coverage over 5’, 3’ UTRs and CDS as reads per kilobase per million mapped in Excel. We used custom shell scripts (makeLengthDistrosOverFeatures.sh, makeLengthDistrosOverNonTrans.sh) to compute the length distributions of the aligned reads from the lengths of aligned sequences.

#### Analysis of P-site periodicity with RiboWaltz

We used the R-package RiboWaltz[38] to infer P-sites by first inferring the optimal offsets from the 5’ and 3’ ends of reads to the start codon, then using these offset estimates to define P-site coordinates for all reads mapping to coding transcripts, then testing whether these P-site coordinates show 3-nt periodicity and/or preference for the predominant reading frame. RiboWaltz requires alignments to the transcriptome as input, so we used STAR v2.6 to align the non-ribosomal Read 1 reads to the transcripts annotated in Mus_musculus.GRCm38.95.gtf:

STAR --runMode alignReads --runThreadN 8 --outSAMtype BAM SortedByCoordinate --genomeDir $index --readFilesIn $fastq1 -- sjdbGTFfile $gtf --readFilesCommand gunzip –c –outFileNamePrefix $prefix --outSAMmultNmax 1 --quantMode TranscriptomeSAM -- outFilterScoreMinOverLread 0 --outFilterMatchNminOverLread 0 -- outFilterMatchNmin 21.

We next used the RiboWaltz package to perform P-site analysis. We loaded the transcriptomes’ alignment and annotation using the RiboWaltz functions “bamtolist()” and “create_annotation()”. We then filtered for reads between 27:33 using the RiboWaltz “length_filter()” function. Next, we inferred P-site offsets for each sample using the “psite()” function, and estimated P-site coordinates of each read using the “psite_info()” function. To plot heatmaps of P-site signal, we ran the “reads_psite_list()” function, extracted estimates of P-site density at each position, and normalized the signals for depth by dividing the total number of CDS-aligned reads. To plot the percentage of P-sites in each frame, we first tabulated the numbers of reads in each frame with the “frame_psite()” function, then plotted the percentages.

#### Gene detection sensitivity and Ribo-seq sample clustering analysis

We calculated gene-level read counts using the deduplicated alignments and subRead:featureCounts, separately for 3’ UTRs, 5’ UTRs, and CDS. We summed read counts in each sample for each gene across 3’ UTRs, 5’ UTRs, and CDS for alkaline fragmented libraries, but used only CDS read counts for Ribo-seq libraries. We used these per-gene, per-sample read counts to assemble a DESeq2 counts matrix. To calculate translation efficiencies, and to identify genes with significantly skewed T.E., we performed a DESeq2 differential expression analysis comparing the AF and RP libraries:

`sig=0.1`

`dds <- DESeqDataSetFromMatrix(countData = DESeq2_counts_matrix, colData = colData, design=∼exp)`

`dds$exp <- factor(dds$exp, levels=c(“AF”,“RP”))`

`dds <- DESeq(dds)`

`res_TE <- results(dds, contrast=c(“exp”, “RP”,“AF”), alpha=sig)`

Results of this DESeq2 analysis include the values “log2FoldChange”, equivalent to the log RP:AF ratio, and thus equivalent to T.E, and “baseMean”, which is the mean expression value for each gene across all samples (“Transcript-per million mapped”, TPMM normalized), and “padj”, the adjusted p-value threshold. We used these values to generate the MA- and volcano-style plots in Figure 6, performing this analysis using the five paired RP and AF samples generated from single cortical hemispheres, since these samples were all generated at the same developmental stage (P4).

DESeq2 removes genes with insufficient counts to meet its independent filtering cutoff, so we utilized this filtered set of genes to calculate pairwise correlation coefficients between all pairs of experiments. We calculated TPMM values for each gene in each sample, then computed Pearson correlation coefficients between samples using all DESeq2 genes. We produced a heatmap and clustering of samples using pheatmap with default parameters, incorporating all CPN samples (to enable comparison of both the three higher input samples generated from P3 CPN, and the P4 samples generated from single cortical hemispheres across a range of inputs).

#### Identification of translated ORFs with RiboCode

We extracted the transcripts encoded by the nuclear genome in Mus_musculus.GRCm38.95.gtf (creating GTF_nucGenome.gtf), and assigned the sequence names of the mm10_genome to match those of the nuclear chromosomes (creating mm10_genome.fasta) in the nuclear gtf file using custom unix scripts. We next prepared the transcripts for RiboCode analysis by running: `prepare_transcripts -g GTF_nucGenome.gtf -f mm10_genome.fasta - o RiboCode_annot.gtf`.

We used RiboCode to produce “metaplots”, which are used to infer the optimal P-site offsets for reads of a given length based on the distribution of 5’ and 3’ end distances from start and stop codons (very similar to how RiboWaltz infers P-sites). We ran: `metaplots -m 27 -M 36 –a RiboCode_annot.gtf -i CPN.dedup.bams`

to produce metaplots_pre_config.txt (this file contains the pairing of BAM file, read length, and optimal P-site offsets). We pooled together all CPN data sets and ran RiboCode to identify translated ORFs:

`RiboCode –a RiboCode_annot.gtf –c metaplots_pre_config.txt -A CTG,GTG,TTG -l yes -b --min-AA-length 6 -o “RiboCode_ORFs”`.

This produces calls for translated ORFs both within each sample, and within a “collapsed” set of translated ORFs, which is the union of all translated ORFs across all samples. We took this “collapsed” data set, and used it to analyze ORF length and start codon distributions using a custom R-script. To plot ORF density across a transcript, we used RiboCode to make the plot:

`plot_orf_density –a RiboCode_annot.gtf -c metaplots_pre_config.txt -t $txID -s $ORFstart -e $ORFend –o $outName`

in which $txID is the transcript ID, $ORFstart and $ORFend are the start and end coordinates of the predicted ORF (with respect to the transcript), and $outName is the output file.

To make the inset plots of the total number of P-sites in each reading frame for an ORF, we obtained the positions of all P-sites inferred by RiboWaltz over the ORF, assigned them to a reading frame, and summed P-site counts in each reading frame using a custom R-script.

#### TopGo Gene Ontology Analysis of uORF-containing transcripts

We used the TopGo R-package to perform Gene Ontology analysis of biological processes, comparing the observed number of uORF containing transcripts to the number expected from a background derived from the “collapsed” set of annotated protein coding transcripts identified by RiboCode across all samples. We used Fisher’s exact test as the test statistic, and plotted p-values of Top-10 most significant ontologies.

---

**Figure 1-figure supplement 1:**
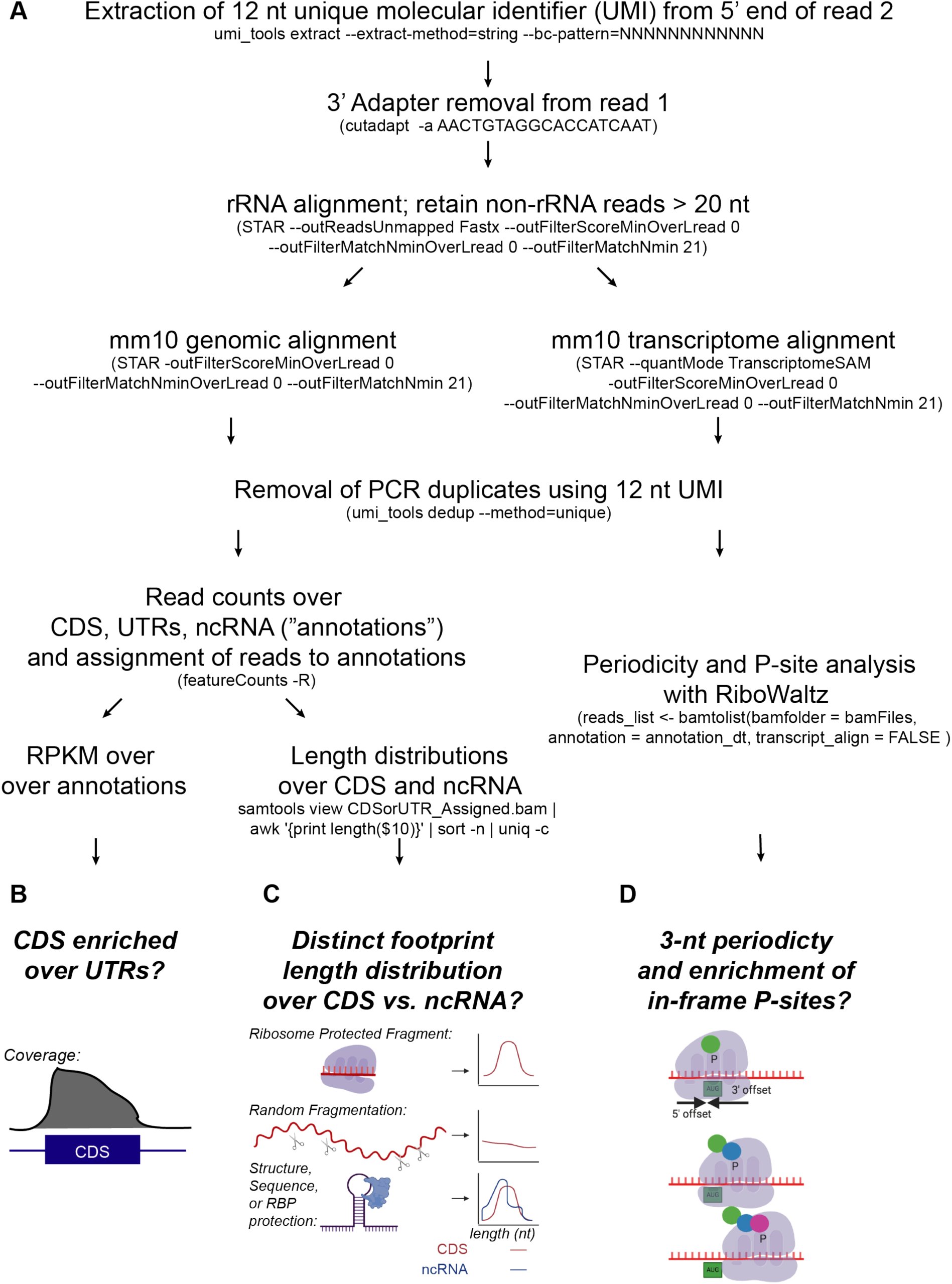
Alignment and quality control for nanoRibo-seq. A) Schematic of alignment and quality control workflow. We first extracted the 12-nt UMI on Read 2. Next, we adapter trimmed Read 1 using the Cutadapt package, then aligned to rRNA with STAR. The reads that do not map to rRNA were aligned to the mm10 genome and transcriptome with STAR. We obtained read counts of reads mapping to 5’ UTRs, CDS, and 3’ UTRs using subRead:featureCounts and a Unix script; read length distributions were generated from BAMfiles using Samtools and a Unix script. This allowed us to evaluate two quality control metrics: enrichment for CDS (B), and comparing length distributions between CDS and various non-coding RNAs (C). We used the R package RiboWaltz[38] to infer P-sites using the observed offsets between the 5’ and 3’ ends of reads covering start codons from the transcriptomes alignments. We plotted P-site heatmaps as a function of distance from start and stop codons, and calculated the percentage of P-sites in each reading frame, a crucial quality control metric (D).

**Figure 2-figure supplement 1:**
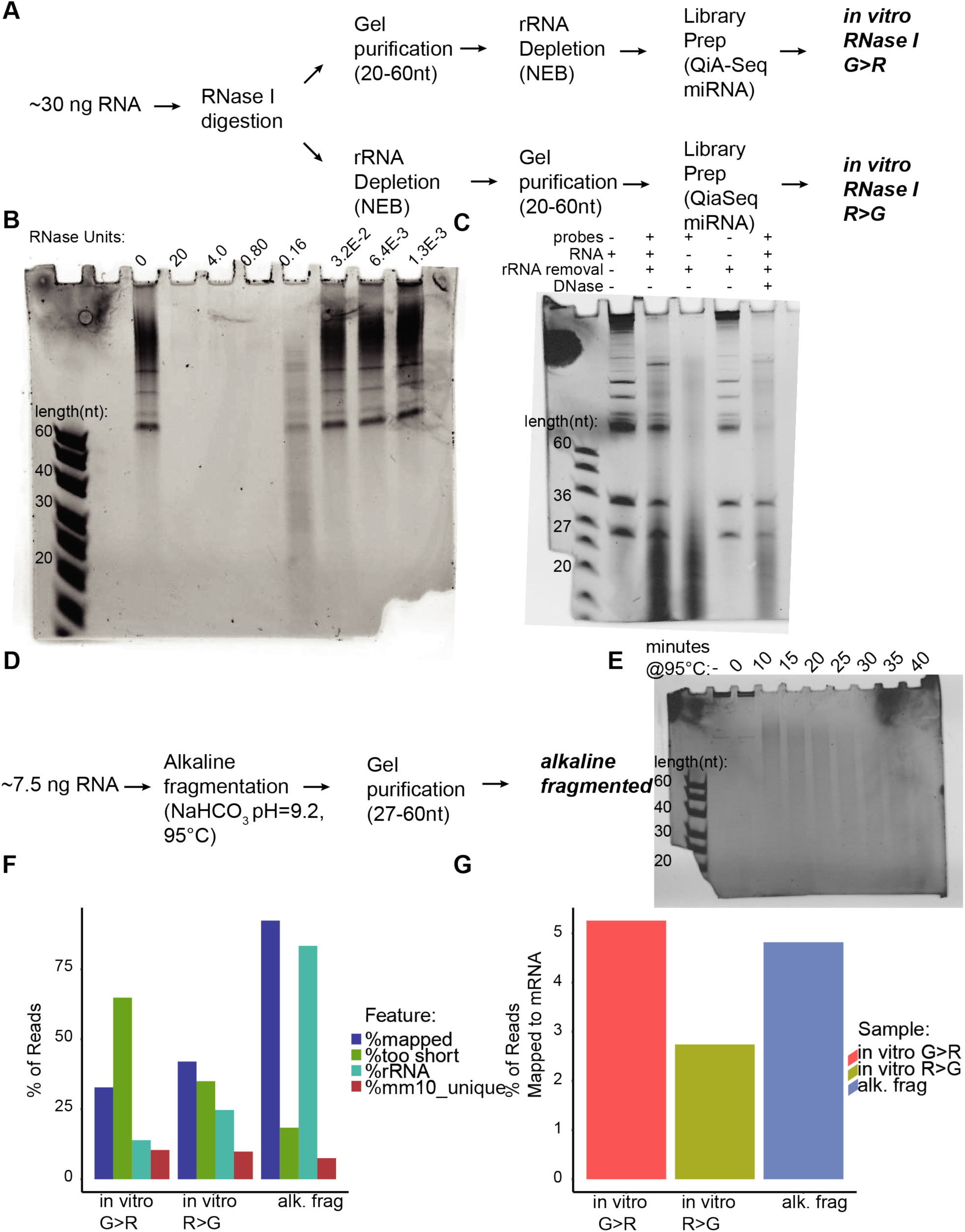
Optimization of total RNA fragmentation with very small quantities of RNA favors alkaline fragmentation. A) Workflows for fragmentation with RNase I. We considered two strategies for RNase I fragmented libraries with minimal rRNA contamination, and compatible with the QiA-seq miRNA kit used to produce RPF libraries: 1) gel purification of 20 to 60 nt RNA before rRNA depletion (G>R method) 2) rRNA depletion before gel purification (R>G method). B) RNase I titration against 30 ng purified brain lysate RNA. C) rRNA depletion probes run in the approximate size range of fragments of interest, and are not fully removed by short DNase digestion (note: 27 and 36 nt standards were added to all lanes to check small RNA degradation during the hybridization and RNase H steps in the rRNA removal protocol). Lane order: 1. RNA, no treatment; 2. RNA with NEB rRNA depletion protocol with probes; 3. rRNA depletion protocol, no RNA added (“probe only” control); 4. RNA with NEB rRNA depletion protocol, no probes added; 5. RNA with rRNA depletion protocol, and with DNase treatment. Note the large amount of material <30 nt in all conditions with probe. D) Workflow for alkaline fragmentation. E) Alkaline fragmentation time titration against 7.5 ng purified cortical lysate RNA. F) % reads mapped to rRNA or mm10 (blue); too short for alignment (<21 nt; green); mapped to rRNA (turquoise); uniquely aligned to mm10 (red) in the three total RNA fragmentation conditions. G) % reads uniquely aligning to mRNA in the three total RNA fragmentation conditions.

**Figure 4-figure supplement 1:**
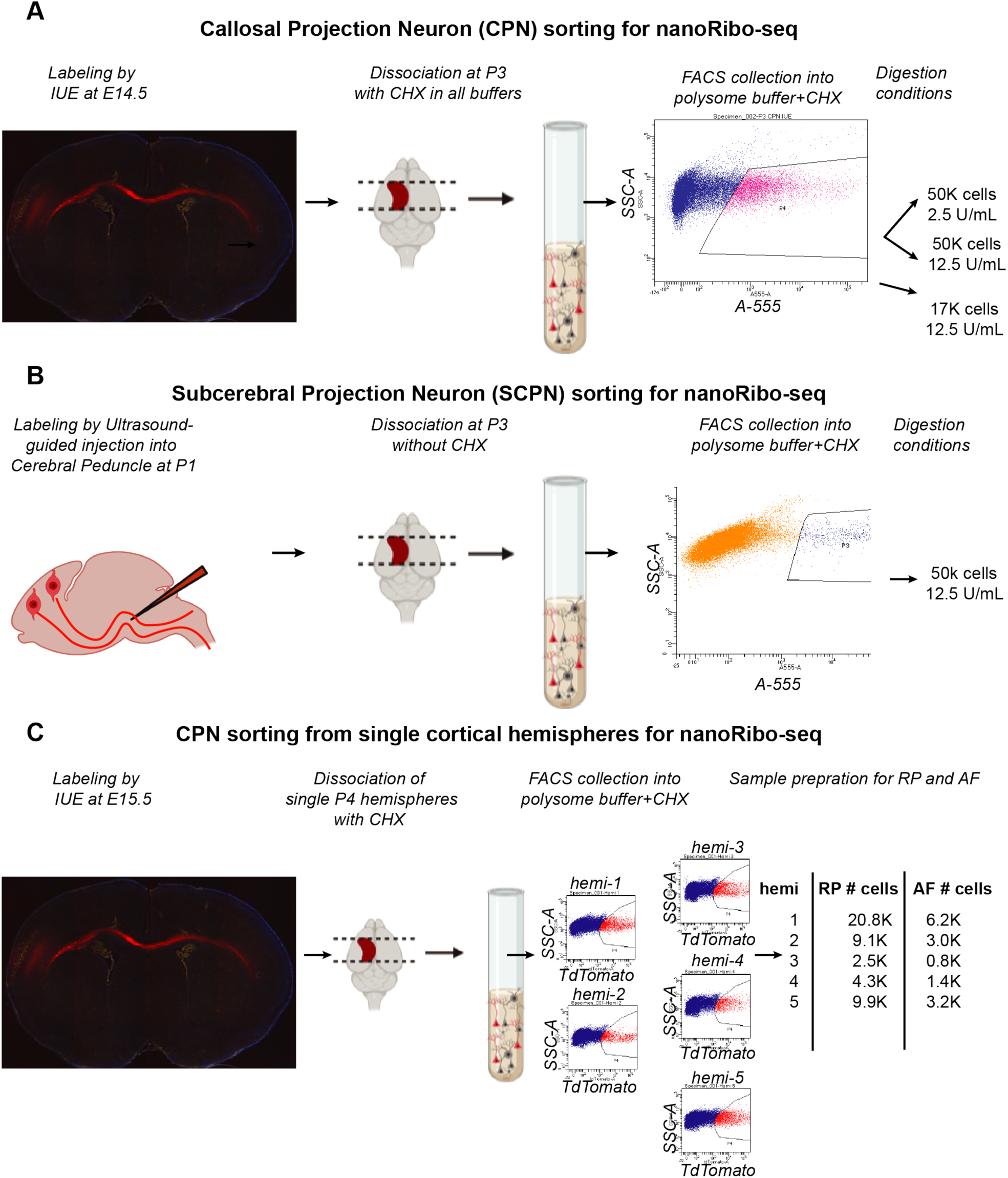
Neuronal subtype labeling, FACS purification, translational inhibition, and RNase I digestion in callosal projection neurons (CPN) and subcerebral projection neurons (SCPN) A) CPN workflow: We labeled CPN by IUE at E14.5. We dissected and dissociated labeled cortices at P3 (in the presence of cycloheximide from tissue isolation onward). Dissociated neurons underwent FACs to purify fluorescent neurons into a polysome lysis buffer containing cycloheximide. We performed three experiments: two libraries generated from 50K CPN on the same day with different [RNase I], and one library generated from a separate sort using 17K CPN. B) SCPN workflow: We retrogradely labeled SCPN by A555-CTB injection into the cerebral peduncle at P1. We dissected and dissociated labeled cortices at P3 (in the absence of cycloheximide). Dissociated neurons underwent FACS to purify 50K fluorescent neurons into polysome lysis buffer containing cycloheximide. C) Workflow for nanoRibo-seq from CPN purified from single cortical hemispheres. CPN were labeled by IUE at E15.5, then prepared for FACS at P4 using individual cortical hemispheres for each sample. CPN were sorted into polysome collection buffer, and split for both either Ribo-seq or RNA extraction and alkaline fragmented.

**Figure S7-figure supplement 1:**
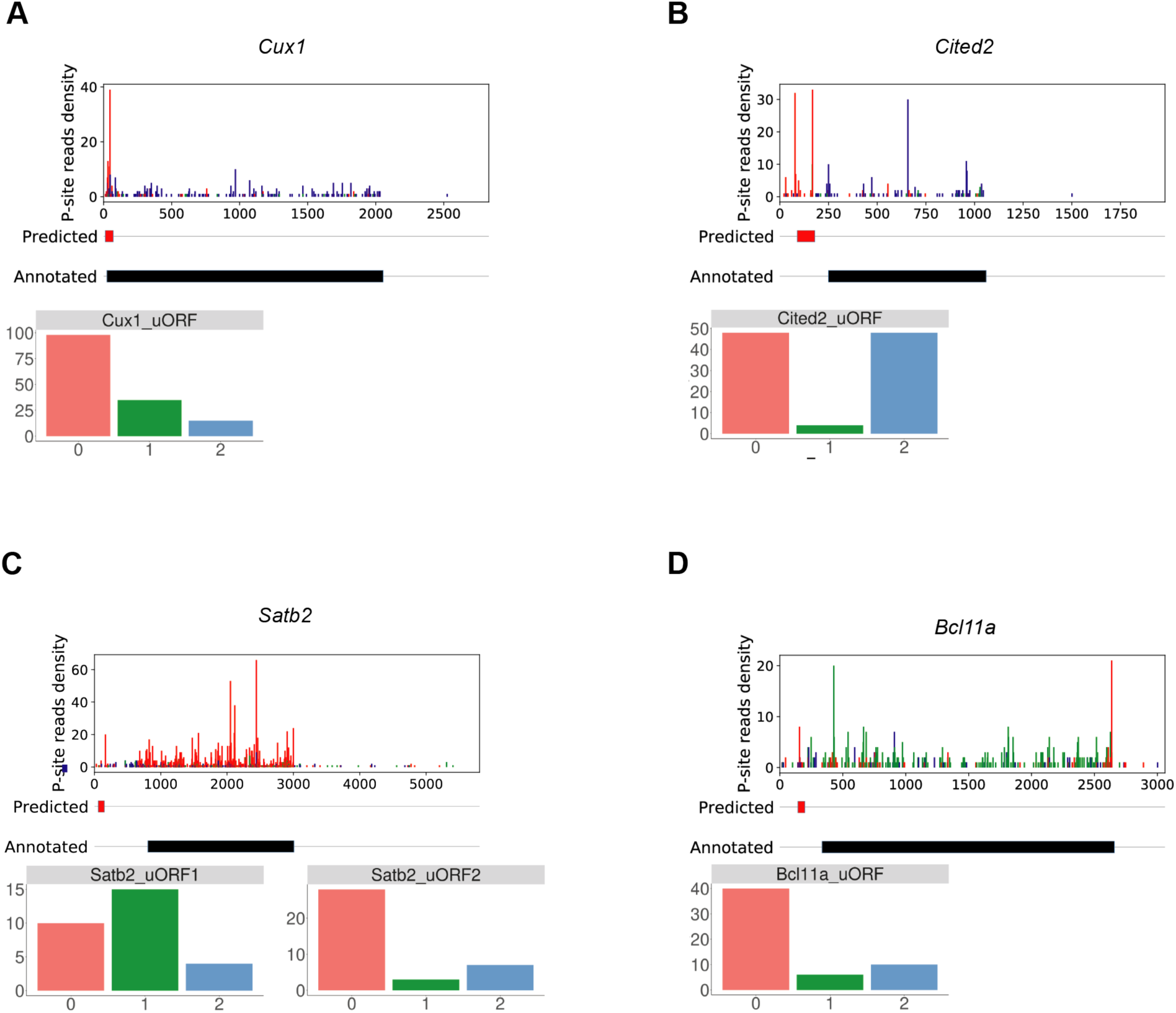
Examples of translated uORFs in the 5’UTRs of TFs important for CPN identity. P-site density plots and number of P-sites in each frame for superficial cortical layer CPN transcription factors A) Cux1, B) Cited2, C) Satb2, and D) Bcl11a. The colored boxes below the density plot shows the position of the predicted uORFs (the first uORF frame is red; other uORFs in the same gene are red if they are in the same frame, blue if they are +1, or green if they are +2 when compared to the first uORF). The black box shows the annotated CDS (regions before and after are 5’ and 3’ UTRs, respectively). The boxes to the right depict the number of P-sites in reading frame: 0 (in-frame, red), +1 (green), +2 (blue), and applies for all panels.

**Figure 7-figure supplement 2:**
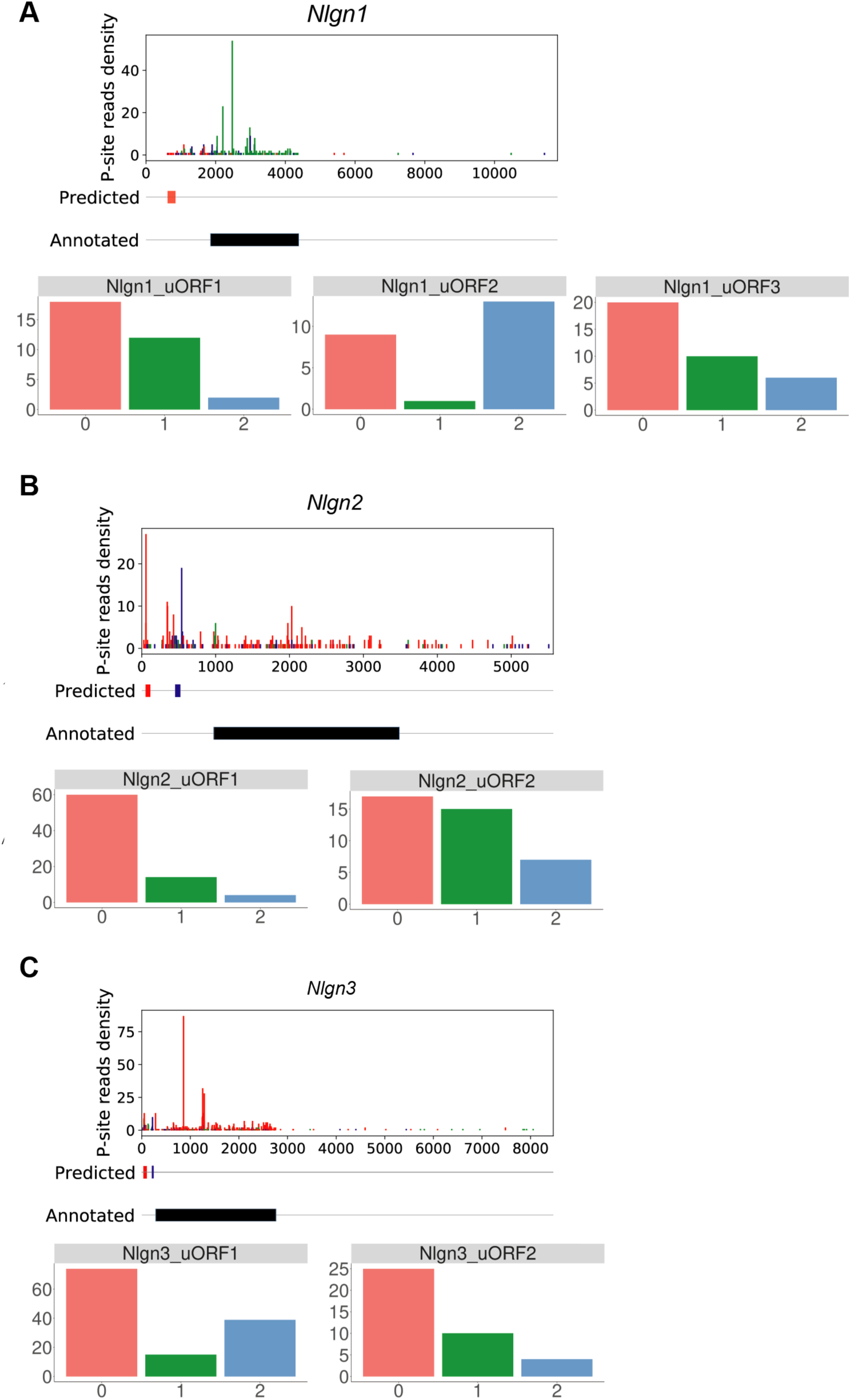
Examples of translated uORFs in the 5’UTRs of neuroligin 1, 2, and 3. P-site density plots and the number of P-sites in each frame for neuroligins Nlgn1, 2, and 3 (A-C, respectively).

**Figure 7-figure supplement 3:**
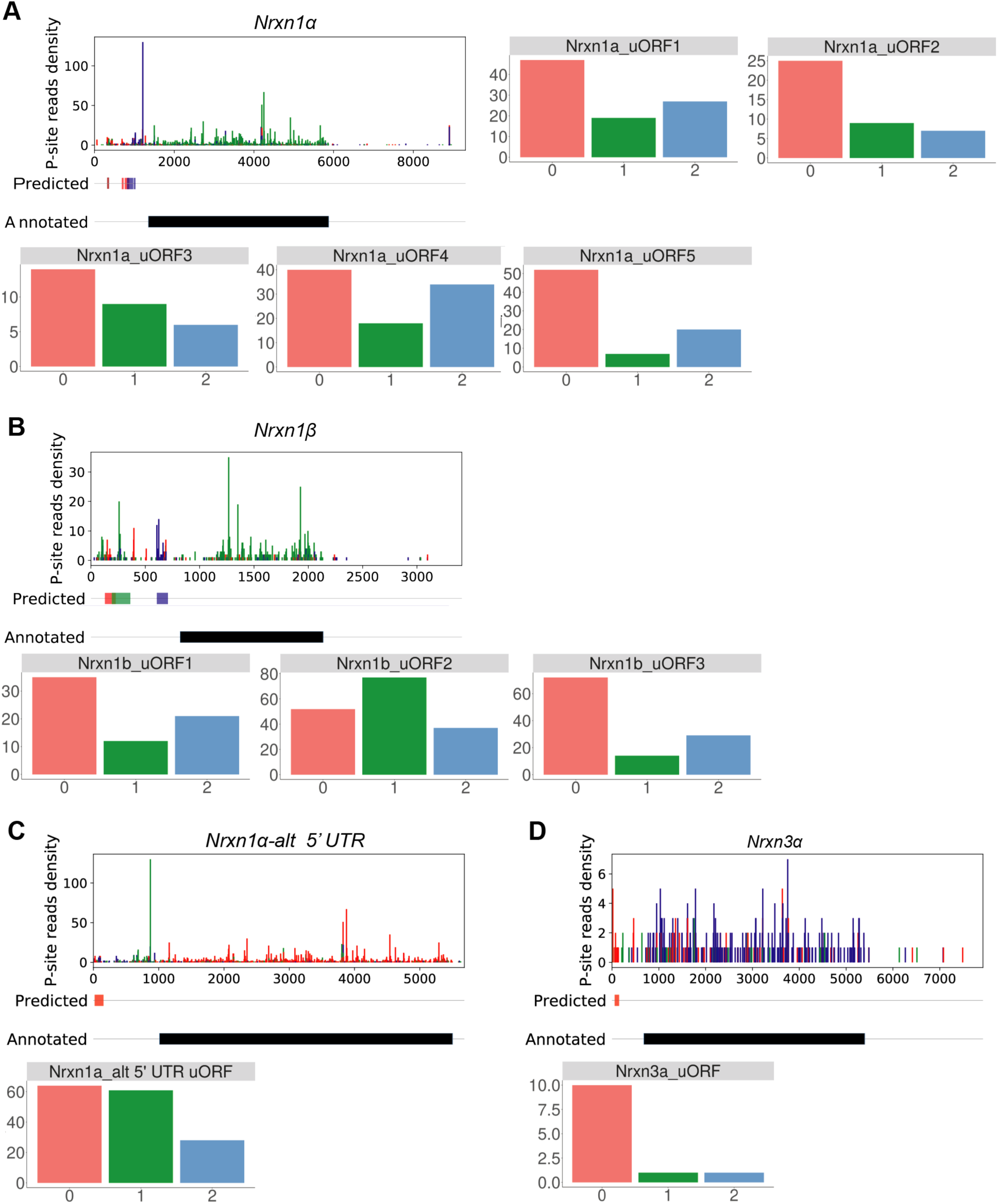
Examples of translated uORFs in the 5’UTRs of Nrxn1 and Nrxn3. P-site density plots and the number of P-sites in each frame for uORFs within the (A) Nrxn1α (transcript ID: ENSMUST00000160844); (B) Nrxn1β, (C) Nrxn1α-alt TSS (transcript ID: ENSMUST00000160800), and (D) Nrxn3α 5’UTRs.

**Figure 7-figure supplement 4:**
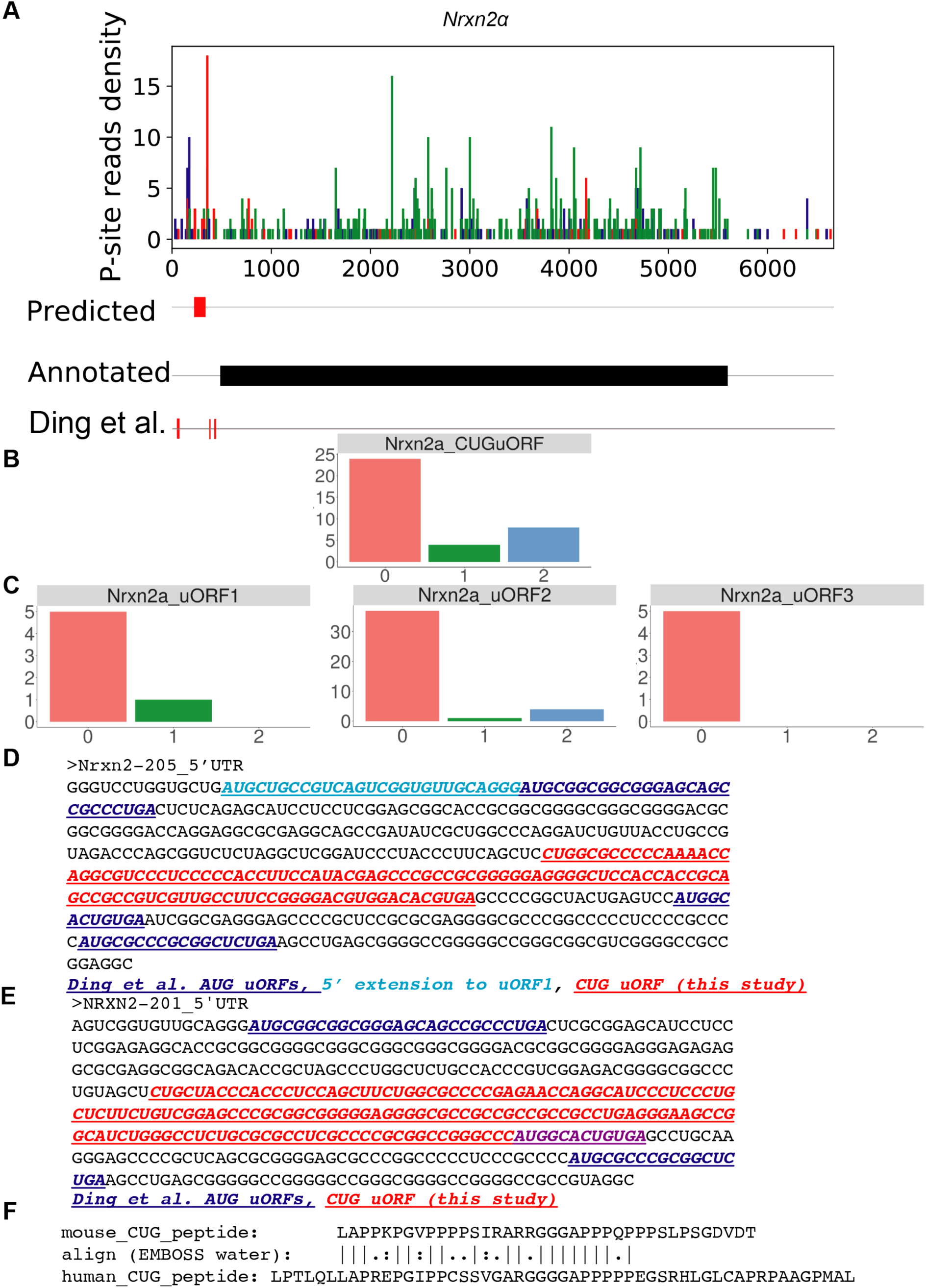
Nrxn2α 5’ UTR contains short, conserved, translated uORFs and a novel translated CUG-start uORF encoding a proline-rich polypeptide sequence. A) P-site density plots and number of P-sites in each frame for the predicted CUG uORF (top) identified in this study, and for the three short AUG ORFs identified by Ding et al. 2020 [52] (uORFs1-3, bottom). Of note, the reference Nrxn2 transcript sequence contains a 5’ extension not analyzed in Ding et al., which contains a possible alternate start codon in-frame with uORF1. B) Mouse Nrxn2α 5’UTR sequence. Red: CUG uORF identified in this study. Blue: short AUG uORFs previously analyzed in Ding et al. Light blue: a 5’ in-frame extension to uORF1 not analyzed in Ding et al., but present in the annotated Nrxn2α 5’ UTR sequence. C) Number of P-sites in each frame for short AUG uORFs. C) Human Nrxn2α 5’UTR sequence. Red: CUG uORF. Blue: short AUG uORFs previously analyzed in Ding et al. Purple: Overlap between CUG uORF and Ding et al. uORF2. D) EMBOSS Water alignment of mouse and human Nrxn2α CUG uORF peptides.

## Notes

### Competing Interest Statement

The authors have declared no competing interest.

